# Modular presynaptic assemblages scale to postsynaptic partner number

**DOI:** 10.64898/2026.01.07.698224

**Authors:** Vanessa M. Puñal, Emma M. Thornton-Kolbe, Jasmine Dhillon, Jackson A. Rogow, E. Josephine Clowney

**Author notes:** Present Address: Department of Neuroscience, Albert Einstein College of Medicine, Bronx, New York, USA.

## Abstract

Behavioral diversification can arise through, and is constrained by, evolutionary and inter-individual differences in neural circuit development. Moreover, alteration of focal neural parameters changes the environment in which cells connect into circuits. In the mushroom body, an associative learning center of arthropods, the number of principal Kenyon cells varies widely across species and among individuals. How such variation is developmentally accommodated by projection neurons, which provide sensory input to Kenyon cells, is not understood. In *Drosophila melanogaster*, we previously demonstrated that projection neurons scale their presynaptic bouton number to Kenyon cell population size. Here, we identify the developmental mechanisms underlying this input flexibility. Boutons arise from projection neuron axonal collaterals; we find that a PN’s collateral number is subtype-specific and serves as the substrate through which bouton number scales to Kenyon cell population size. Independent of projection neuron identity or Kenyon cell number, individual collaterals most often produce just one bouton, suggesting collaterals are modular cell biological bouton units. Developing projection neurons initially overproduce nascent collaterals in the early pupa. The set of nascent collaterals that mature and eventually bear boutons is conditional on Kenyon cell number, thereby executing scaling. Finally, early boutons bear filopodia that frequently contact neighboring PN processes, suggesting that bouton–bouton interactions contribute to shaping these structures.

## Introduction

Changes in neural circuit development diversify behavioral repertoires over evolutionary time. One mechanism of change is via alterations in cell number. Increases in the absolute number of neurons in associative brain centers across mammalian species is correlated with the acquisition of flexible, complex cognition (Herculano-Houzel, 2017; Ströckens et al., 2022). Interestingly, within species, the precise numbers of neurons in associative and other brain centers can also vary substantially, suggesting flexibility in developmental programs (Azevedo et al., 2009; Herculano-Houzel et al., 2015; Lu et al., 2001; Neves et al., 2020; Strom and Williams, 1998). While the mechanisms that expand cell number are well understood, how developmental programs that pattern connectivity accommodate changes in the ratio between pre- and postsynaptic partners is less clear. In columnar structures such as the cortex or visual system, pre- and postsynaptic neurons often arise from a shared pool of neuroblasts (Gao et al., 2014). As a result, changing the number of progenitors proportionally scales both circuit layers. In contrast, other neural circuits are constructed of pre- and post-synaptic layers generated by distinct neurogenic programs (Ito et al., 2013), where altering neuron number in one layer alone disrupts the natural ratio of pre- to postsynaptic cells. Understanding the developmental mechanisms that enable neurons to reliably accommodate differences in partner number and produce circuits capable of driving behavior could offer insight into how these mechanisms enable and constrain change over evolutionary time.

Here, we use an associative learning center of arthropods, the mushroom body, to explore such mechanisms. Across insect taxa, the number of principal neurons in each mushroom body hemisphere, called Kenyon cells (KCs), varies widely: from 2,000 in *Drosophila melanogaster,* to between 10,000 and 80,000 in Heliconiini butterflies, and over 300,000 in the beetle *Cotinus mutabilis* (Couto et al., 2023; Farris, 2013; Li et al., 2020). Evolutionary expansion of KC number is correlated with differences in species-specific learning capacity (Ellis et al., 2024; Farnworth and Montgomery, 2024; Farnworth et al., 2024; Young et al., 2024). KC number can also differ substantially between conspecifics, and is influenced by environmental conditions during development (Heisenberg et al., 1995; Lin et al., 2013; Schlegel et al., 2024). Despite taxonomic and individual-to-individual differences in KC number, development consistently produces functional circuits. Indeed, we have found that flies retain olfactory associative learning ability even when KC numbers are engineered to be ¼ to 2x normal population size (Ahmed et al., 2023). In other contexts in the *D. mel* nervous system, trophic cell death has been shown to match the ratio of pre- and post-synaptic cells, yet we have not observed cell death in mushroom body development (Courgeon and Desplan, 2019; Elkahlah et al., 2020).

KCs require multiple active inputs to fire, and thus behave as coincidence detectors to encode odor identity (Gruntman and Turner, 2013). Each olfactory KC produces 3-10 dendritic claws that connect to 3-10 PN presynaptic axonal boutons from among 52 types of olfactory projection neurons (Aso et al., 2009; Bates et al., 2020; Jefferis et al., 2001; Stocker et al., 1997). This sparse, combinatorial connectivity maximizes odor representation: too many inputs onto individual KCs saturates KC population responses to odors, while too few inputs reduces the number of odors the KC collective can represent (Ahmed et al., 2023). In previous work, we manipulated the number of Kenyon cell neuroblasts during development to alter KC population size and found that presynaptic PN bouton numbers scale to the KC population (Ahmed et al., 2023; Elkahlah et al., 2020). In contrast, KC dendrite number remained stable in this context, as well as in the face of developmental changes to PN population size (Ahmed et al., 2023; Elkahlah et al., 2020). The plasticity of PN morphogenesis and rigidity of KC dendritic production together ensure maintenance of adult KC population-level odor responses when KC numbers change (Ahmed et al., 2023; Elkahlah et al., 2020).

Though the total number of PN boutons is developmentally plastic, individual cells of each of the 52 uniglomerular PN types bear characteristic numbers of boutons, which are located in predictable, subtype-specific locations within the adult calyx (Jefferis et al., 2007; Lin et al., 2007; Marin et al., 2002; Tanaka et al., 2004; Thornton-Kolbe et al., 2025; Wong et al., 2002; Zheng et al., 2018; Zheng et al., 2022). In each hemisphere, stereotyped numbers of PNs are born in a reproducible order in the embryo and larva from two neuroblasts, each of which generates distinct subsets of PNs (Grabe et al., 2016; Lin et al., 2012; Yu et al., 2010). By the onset of pupation, both PN neuroblasts have ceased generating new PNs, and the main axon tract of all embryonic- and larval-born PNs projects through the mushroom body calyx (Jefferis et al., 2004; Marin et al., 2005). In contrast, KCs are born from 4 neuroblasts that divide continuously from embryonic stages until just before adult eclosion (Ito and Hotta, 1992; Prokop and Technau, 1991; Truman and Bate, 1988). Remarkably, at the start of pupation, ∼40% of KCs still have not been born (Aso et al., 2009; Lee et al., 1999). How PNs simultaneously scale bouton production to a KC population whose final size is not yet known, while also maintaining subtype-specific morphological programs, is not understood.

Here, we characterized PN morphological development both in normal calyces and those in which we modified KC number. First, we observed that developing PNs overproduce axonal filopodia just hours after pupation. Only a subset of these filopodia go on to become bouton-bearing collaterals, while the rest are eliminated. Second, we observed across PN subtypes from calyces with normal or altered KC population size that variation in bouton number was predicted by the number of axonal collaterals produced during development. Individual collaterals most often hosted just one bouton. We consistently found this pattern across different PN subtypes both in EM connectomes as well as *in vivo* over engineered KC population sizes. The concordance of collateral number with bouton number and the narrow range of the number of boutons per collateral suggested that PNs quantitatively matched KC population by adjusting collateral stabilization. Indeed, we found that PN collateral production scales to KC population size as soon as PNs begin sprouting axonal filopodia in the very early pupa. In the complete absence of post-embryonic KCs, newly sprouted axonal filopodia display a reduced capacity for outgrowth and are eventually lost. Strikingly, all PNs have been born well before this point in development, but KC neurogenesis will continue for several days, suggesting that PNs stabilize collaterals based on an early correlate of future, final KC number.

## Results

### Projection neurons of different types scale boutons and collaterals to Kenyon cell population size

In the adult mushroom body, olfactory information from 52 distinct odor channels is relayed to KCs via 52 subtypes of uniglomerular, cholinergic olfactory PNs. The 100 PNs across these 52 subtypes are born from two neuroblasts (NBs): an anterodorsal NB (ALad1, NotchOFF “B” hemilineage) and a lateral NB (ALl1, NotchOFF “B” hemilineage) (Figure 1A). ALad1 generates 18 embryonic, anterodorsal PN types (adPNs), which innervate the larval calyx (Lai et al., 2008; Marin et al., 2005; Masuda-Nakagawa et al., 2005; Yu et al., 2010). Both ALad1 and ALl1 generate additional PN types after larval hatching (22 and 12 types, respectively), which will participate in the adult calyx in conjunction with the embryonically born adPNs (Bates et al., 2020; Jefferis et al., 2004; Lin et al., 2012; Marin et al., 2005; Yu et al., 2010). At metamorphosis, embryonic adPN calyx innervations used in the larva are pruned away; then, during early pupal timepoints, all uniglomerular olfactory PNs initiate new neurite production towards construction of an adult calyx (Marin et al., 2005; Puñal et al., 2021).

**Fig 1.**
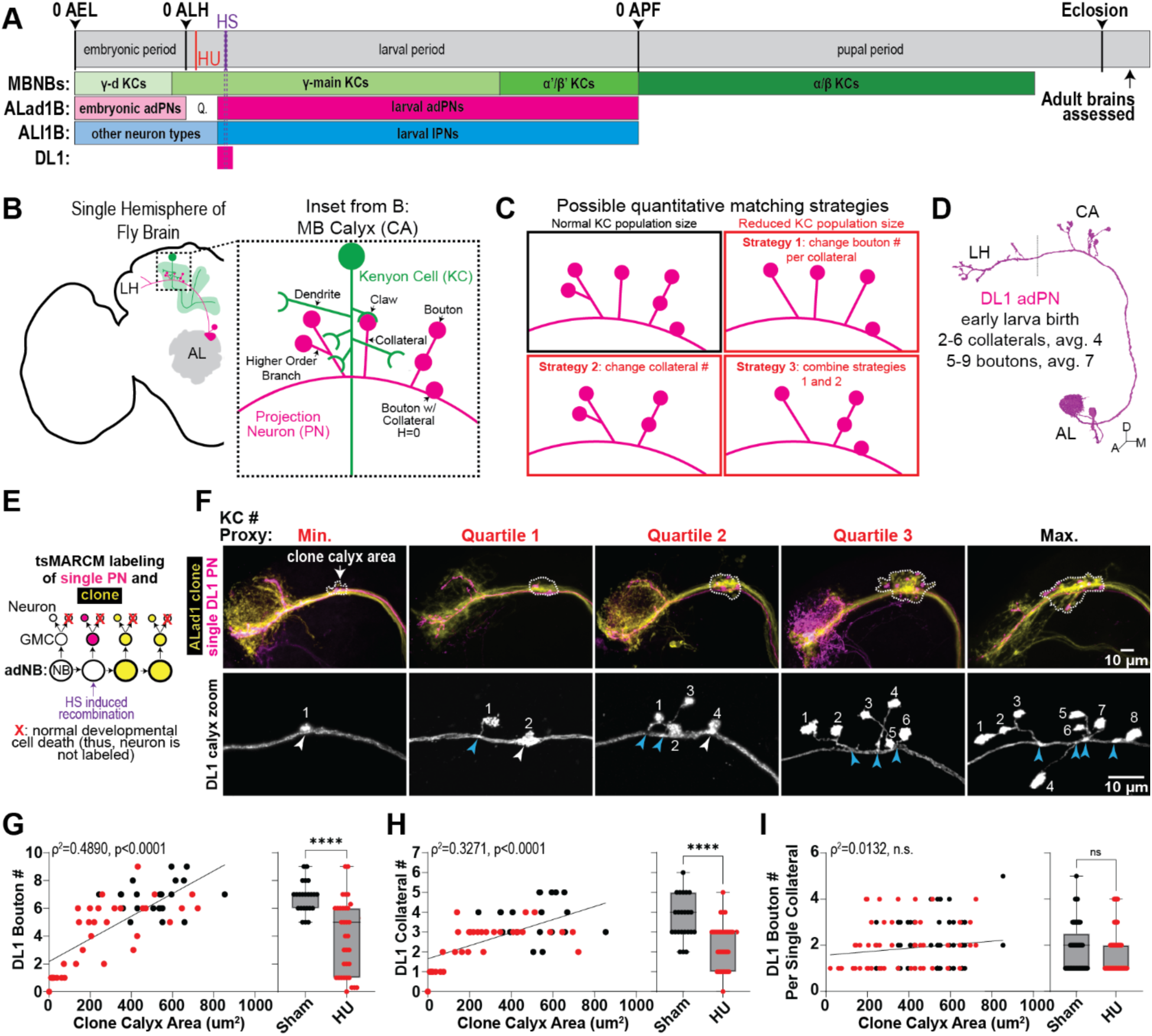
Early larval-born DL1 adPNs scale collaterals to KC population size. **A:** Timeline for neuronal birth. Hydroxy-urea ablation (HU, red line) and heatshock (HS, purple line) timepoints for induction of twin-spot MARCM (tsMARCM) labeling annotated. Q: Quiescence. AEL: after egg laying. ALH: after larval hatching. APF: after puparium formation. MBNB: mushroom body neuroblast lineage. ALad1B: anterodorsal PN hemilineage. ALl1B: lateral PN hemilineage. **B:** Schematic of the adult fly brain and olfactory system. AL: antennal lobe. MB: mushroom body. LH: lateral horn. Inset: Cartoon of enlarged MB calyx, depicting the typical morphology and connectivity pattern of KCs and PNs. H: height. **C:** Predicted PN strategies to scale bouton numbers to KC population size. **D:** FlyWire reconstruction of a single adult DL1 PN. Dashed line demarcates the boundary between the MB calyx (CA) and lateral horn (LH). **E:** Schematic of genetic strategy utilized to label single adult DL1 adPNs and clone of subsequently born adPNs. Magenta: single PN labeling. Yellow: clonal labeling of PNs born after HS. Red X: dead neuron, which is an end-product of normal development in the adNB lineage (Lin et al., 2010). GMC: ganglion mother cell. NB: neuroblast. **F:** Adult *GH146*+ tsMARCM clones (top: yellow) and single DL1 PNs (top: magenta, bottom: white) across varying KC population sizes induced via HU ablation (red titles), or sham treatment as control (black title). White dashed lines (top row) outline the calyx area defined by boutons of the clone, which serves as a proxy for KC number. Bottom row: White arrowhead: collateral of height zero. Light blue arrowhead: collateral of non-zero height. Numbers indicate bouton counts. **G-H:** Scatter plot with simple linear regression lines (left) and box-and-whisker plot (right) showing the relationship between DL1 bouton number (G) or collateral number (H) and clone calyx area. Each point represents a single neuron (black: sham; red: HU). Statistics: two-tailed Spearman’s correlation (scatter plots) or two-tailed Mann-Whitney test (box-and-whisker plots). \*\*\*\**p* < 0.0001. **I:** (Left) Scatter plot with simple linear regression lines (left) and box-and-whisker plot (right) showing the relationship between DL1 bouton number per single collateral and clone calyx area. Each point represents a single collateral (black: sham; red: HU). Statistics: two-tailed Spearman’s correlation (scatter plot) or two-tailed Mann-Whitney test (box-and-whisker plots). n.s.=0.1732 (scatter plot), 0.3805 (box-and-whisker plot).

In the adult calyx, PNs bear agglomerations of presynaptic sites called boutons which KC dendrites enwrap, forming a structure called a microglomerulus. These boutons are organized in the calyx at the ends of collaterals which branch off the main axon (Figure 1B). Each PN type has a unique and type-specific axonal morphology and places boutons in predictable locations across the calyx (Thornton-Kolbe et al., 2025). We observed in our previous work that adult PN bouton number is flexible and scales to developmental manipulations of KC number (Elkahlah et al., 2020). Bouton scaling to KC population size could be the outcome of PNs developmentally altering: 1) bouton number per collateral while keeping total collateral number stable, 2) total collateral number with no alterations to bouton number per collateral, or 3) altering both collateral number and bouton number per collateral (Figure 1C). As a population, all PNs could use just one of these strategies to scale bouton numbers or, given that PN subtypes are transcriptionally different from each other throughout pupal development and morphologically unique in the adult, PNs could use type-specific strategies (Li et al., 2017; Xie et al., 2021). To elucidate the morphological approach(es) taken by PNs, we assayed the adult morphologies of specific PNs from animals with experimentally imposed variations in KC number during development.

We initially assessed DL1 PNs, the first PN type born in the early larva of the ALad1 lineage (Figure 1A,D). Using twin-spot MARCM (tsMARCM) to visualize birth-timed PNs within the *GH146-GAL4* expression pattern, we induced heat shock at the time of DL1 genesis (Yu et al., 2009) (Figure 1A,E). In adult wildtype brains, we found that single DL1 PNs on average have 4 collaterals and 7 boutons in the calyx (Figure 1G,H). To test how DL1 PNs adjust their development to the KC population size, we combined tsMARCM with either mild hydroxyurea (HU) or sham treatment of newly hatched larvae (Figure 1A,E). HU sporadically ablates KC NBs, as we and others have described previously (Ahmed et al., 2023; De Belle and Heisenberg, 1994; Elkahlah et al., 2020; Sweeney et al., 2012). In addition to labeling DL1 PNs in one color, tsMARCM in the context of *GH146-GAL4* causes the 21 ALad1 lineage adPN subtypes subsequently born after DL1 to be clonally labeled in a second color (Figure 1E,F). Thus, we measured the total calyx area occupied by tsMARCM clone boutons as a proxy for KC population size (Figure 1F), similar measures of which (i.e. PN ChAT+ calyx area) we have previously found to scale with KC number (Ahmed et al., 2023; Elkahlah et al., 2020). At maximal NB ablation, the only KCs remaining are a set of ∼200 KCs born in the embryo, most of which are of the γ-d visual type (Ganguly et al., 2023; Kunz et al., 2012; Puñal et al., 2021; Technau and Heisenberg, 1982)

We quantified the number of boutons, collaterals, and boutons per collateral across each condition and asked how these scaled to calyx size. These data revealed that adult DL1 PN bouton number was significantly reduced in the HU condition relative to control and scaled with calyx size (Figure 1G), recapitulating what we previously reported for other PN types (Elkahlah et al., 2020). Collateral counts followed these same trends (Figure 1H). However, bouton number per collateral was unchanged and largely ranged between 1 to 3 boutons per collateral independent of ablation status (Figure 1I).

We next re-analyzed a set of images of embryonically-born VM4 and VM6 PNs (Elkahlah et al., 2020) to ask whether PN types developmentally and morphologically distinct from DL1 use the same cellular mechanism to adjust their bouton repertoire to KC population size. Despite the closeness of their birth timing (Yu et al., 2010) and shared lineage, VM4 and VM6 PNs diverge morphologically: earlier-born VM4 PNs have an average of 4 boutons per PN, while the VM6 PN averages 10 (Figure 2B,G). In the absence of post-embryonic KCs, adult VM4 and VM6 PN axons lacked collaterals and boutons (Figure 2C,H). Across partial KC ablation conditions, both PN types scaled their total bouton and collateral numbers to calyx area, whereas the number of boutons per collateral remained constant at ∼1–3 across all conditions (Figure 2D–K; Figure 2—Supplemental 1).

**Fig 2.**
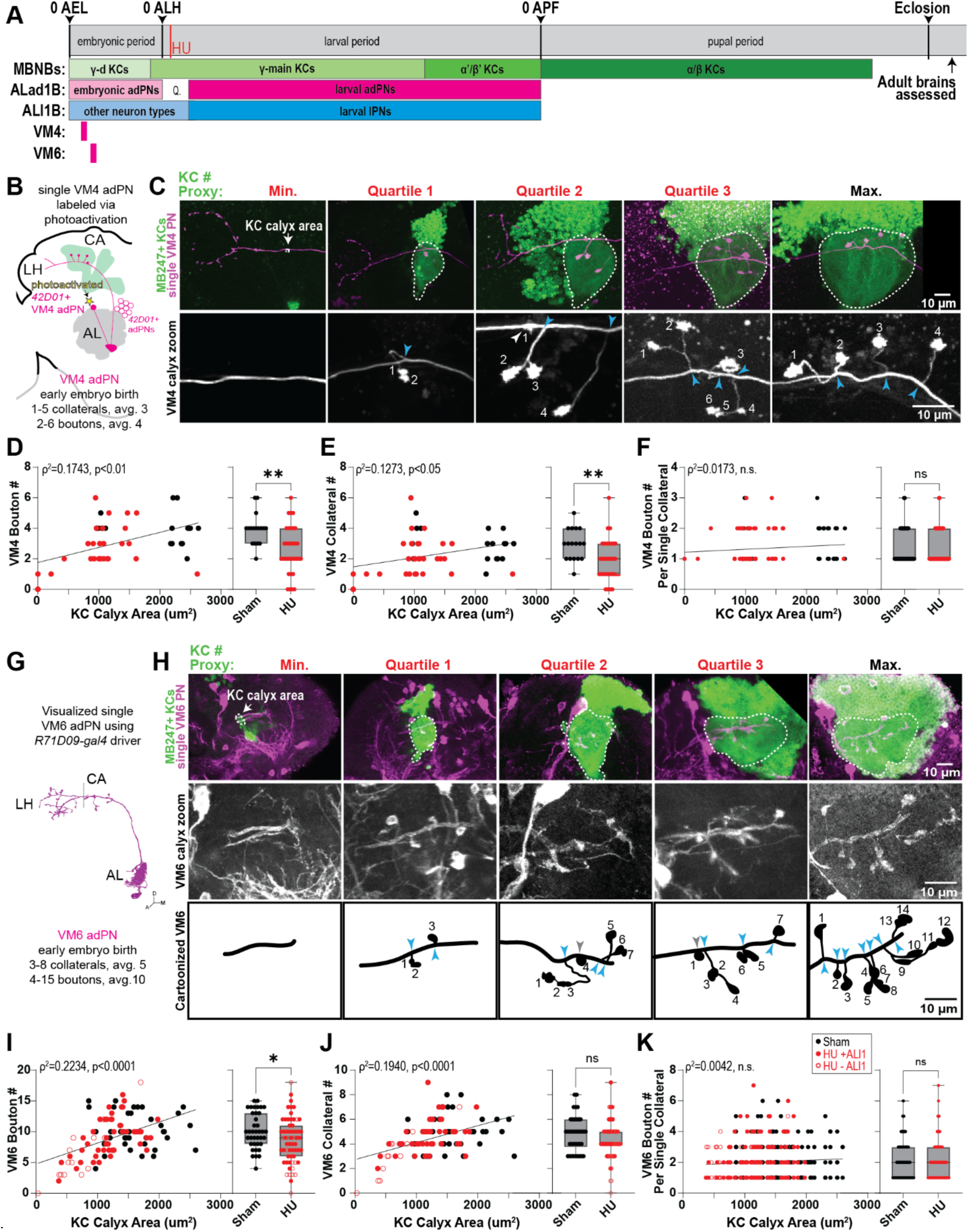
Early embryonically-born VM4 and VM6 adPNs scale collaterals to KC population size. **A:** Identical developmental timeline from Fig 1A. Contextualized here is the birth-timing for early embryonically born VM4 and VM6 adPNs (magenta boxes). **B:** Schematic of one hemisphere of the adult central brain and olfactory system, illustrating strategy utilized to label a single *42D01*+ VM4 adPN. **C:** Adult KCs (green) and single VM4 PNs (top: magenta, bottom: gray) across varying KC population sizes induced via HU ablation (red titles), or sham treatment as control (black title). White dashed line outlines *MB247*+ KC calyx area, which serves as a proxy for KC number. White arrowhead: collateral of height zero. Light blue arrowhead: collateral of non-zero height. Numbers indicate bouton counts. **D-E:** Scatter plot with simple linear regression lines (left) and box-and-whisker plot (right) showing the relationship between bouton number (D) or collateral number (E) for VM4 PNs and KC calyx area. Each point represents a single neuron. (black: sham; red: HU). Statistics: two-tailed Spearman’s correlation (scatter plots) or two-tailed Mann-Whitney test (box-and-whisker plots). \*\**p* < 0.01. **F:** Scatter plot with simple linear regression lines (left) and box-and-whisker plot (right) showing the relationship between VM4 bouton number per single collateral and KC calyx area. Each point represents a single collateral (black: sham; red: HU). Statistics: two-tailed Spearman’s correlation (scatter plot) or two-tailed Mann-Whitney test (box-and-whisker plots). n.s. p-value: 0.1674 (scatter plot), 0.9467 (box-and-whisker plot). **G:** FlyWire reconstruction of a single adult VM6 PN. Dashed line demarcates the boundary between the MB CA and LH. **H:** adult KCs (top: green) and single VM6 PNs (top: magenta, middle: gray) across varying KC population sizes induced via HU ablation (red title), or sham treatment as control (black title). White dashed line outlines *MB247*+ KC calyx area. Manual tracings (bottom) of single VM6 PNs depicted in middle row. Gray arrowhead: collateral of height zero. Light blue arrowhead: collateral of non-zero height. Numbers indicate bouton counts. **I-J:** Scatter plot with simple linear regression line (left) and box-and-whisker plot (right) showing the relationship between bouton number (I) or collateral number (J) for VM6 PNs and KC calyx area. Each point represents a single neuron (black: sham, solid red: +ALl1, open red: −ALl1). Statistics: two-tailed Spearman’s correlation (scatter plots) or two-tailed Mann-Whitney test (box-and-whisker plots). \**p* < 0.05. n.s. *p*-value: 0.1598. **K:** Scatter plot with simple linear regression line (left) and box-and-whisker plot (right) showing the relationship between VM6 bouton number per single collateral and KC calyx area. Each point represents a single collateral (black: sham, solid red: +ALl1, open red: −ALl1). Statistics: two-tailed Spearman’s correlation (scatter plot) or two-tailed Mann-Whitney test (box-and-whisker plot). n.s. p-value: 0.1729 (scatter plot), 0.2683 (box-and-whisker plot).

We note that ALl1 can be sensitive to HU ablation (Elkahlah et al., 2020; Lovick and Hartenstein, 2015). Because loss of other PNs increases bouton production by spared PNs, while loss of KCs decreases bouton production, differences in inter-PN competition across samples could present a confound (Elkahlah et al., 2020; Thornton-Kolbe et al., 2025). However, we found that loss of the ALl1 lineage PNs in addition to KC NB ablation had no effect on VM6 bouton number per collateral across sham and HU conditions (Figure 2 Supplemental 1D). This suggests bouton number per collateral is stable even to changes in PN population size.

### Bouton number scales with collateral production through modular collateral::bouton units

Our data for DL1, VM4, and VM6 suggest that PNs may universally scale bouton numbers to KC population size by adjusting collateral production. To test whether these followed the same mathematical distribution, we compared the regression lines for each individual cell type to that of the whole cohort, and found they had strikingly similar slopes (Figure 3 Supplemental 1A,D,G). These results suggested PNs share a common bouton::collateral relationship independent of KC population size or PN identity.

**Fig 3:**
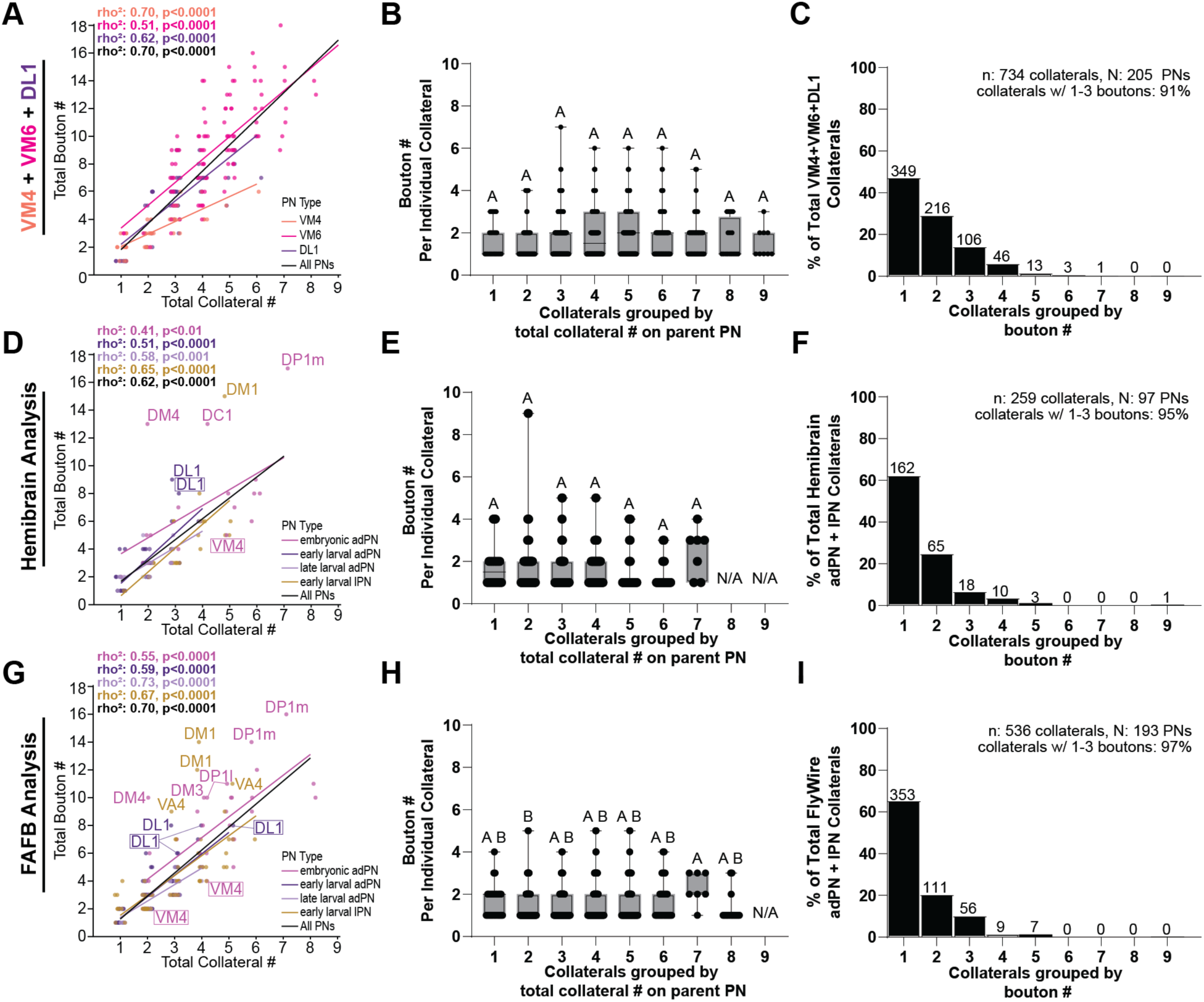
PN collaterals act as modular bouton units. **A–C**: DL1, VM4, and VM6 cells from HU experiments combined into a single cohort. (A) Scatter plot showing the relationship between total collateral number and total bouton number per PN; points colored by PN type. Scatter plots shown with jitter to reduce point overlap. (B) Box-and-whisker plots of bouton number per collateral, with collaterals binned by the total collateral count of the parent PN. (C) Histogram showing the frequency distribution of bouton numbers per collateral. **D–F**: Hemibrain connectome analysis. Same plot types as A-C. In (D), outlier PNs and PN types represented in A–C are labeled; PN names from A-C are outlined with boxes, unless a cell is an outlier. PNs colored by developmental origin. **G–I**: FAFB (FlyWire) connectome analysis. Same plot types as A-C. In (G), outlier PNs and PN types represented in A-C are labeled; PN names from A-C are outlined with boxes, unless a cell is an outlier. PNs colored by developmental origin. Statistics: (Scatter plots) two-tailed Spearman’s correlation; best-fit lines generated by simple linear regression. Each point represents one PN. (Boxplots) one-way ANOVA followed by Kruskal–Wallis, with Compact Display System (CDS) lettering indicating significant group differences (see Figure 3, supplemental table for details). Each point represents one collateral.

To ask whether the linear relationship between bouton and collateral numbers is universal among PN subtypes, we quantified total collateral and bouton numbers for each PN in the Hemibrain (one hemisphere) and FAFB (two hemispheres) connectomes (Dorkenwald et al., 2024; Scheffer et al., 2020; Schlegel et al., 2024; Zheng et al., 2018). Collateral number strongly and significantly predicted bouton number for most PN types independent of developmental origin (Figure 3D,G). A small set of PN types (DP1m, DC1, DM4, DM1) deviated from the wider trend, with DP1m, DM4, and DM1 emerging as outliers—with more boutons per collateral—in both connectomes (Figure 3D,G).

These analyses suggested that, in the natural calyx, a single function relates collateral and bouton number across contexts for most PN types. Therefore, we next quantified the number of boutons found on each individual collateral in each connectomic dataset. We found that the number of boutons any individual collateral made did not vary with the total number of collaterals present on a PN nor its developmental origin (Figure 3B,E,H, Figure 3 Supplemental 1B,E,H, Figure 3 Supplemental 2). >90% of collaterals had three or fewer boutons, and 65% had just one (Figure 3C,F, I, Figure 3 Supplemental 1C,F,I).

Together, these findings indicate different subtypes of PNs in the natural calyx produce subtype-specific numbers of boutons by scaling collateral number, and the same mechanism accounts for their ability to adjust to KC population size. Thus, individual collaterals operate as modular units following a consistent bouton-production function.

### DL1 projection neuron adult morphology is established before Kenyon cell neurogenesis is complete

Our data suggests a universal developmental rule whereby modifications to the number of discrete bouton::collateral assemblages strongly predicts total bouton number across PN subtypes in the natural calyx and across calyces with variable KC number. As collaterals are the backbone of bouton::collateral assemblies, we next sought to determine when adult collateral numbers and morphologies are established in development. To this end, we visualized DL1 adPNs labeled using tsMARCM at select developmental timepoints in wildtype calyces (Figure 4).

**Fig 4:**
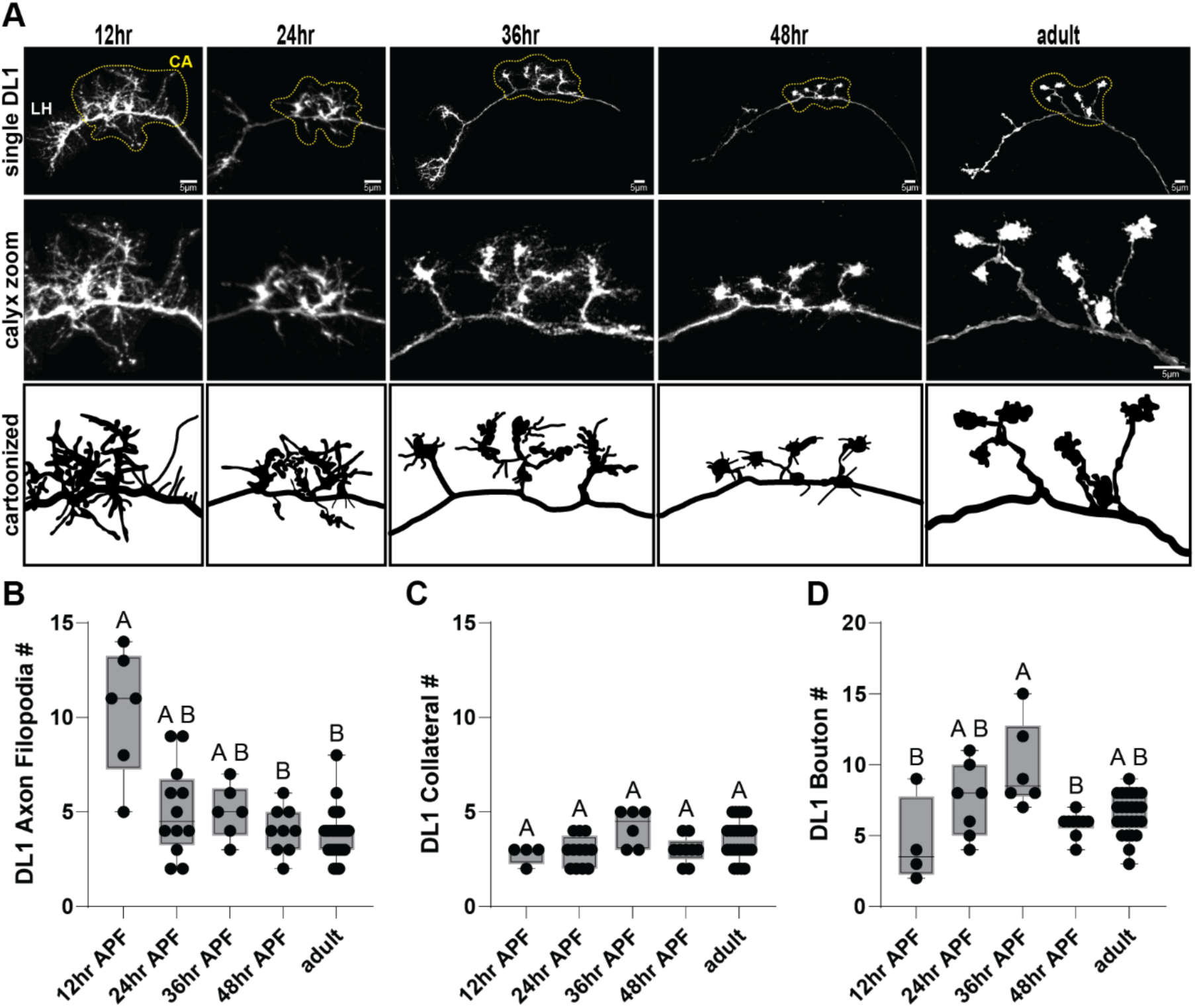
DL1 adPN collateral-bouton assemblages form prior to completion of KC neurogenesis. **A:** Top row: single developing DL1 PNs labeled by twin-spot MARCM. Yellow dashed line outlines the calyx (CA). LH: lateral horn. Middle row: zoomed-in views of the calyx. Bottom row: schematic depictions of the zoomed-in regions at each developmental timepoint. **B–D:** Box-and-whisker plots showing the number of axonal filopodia (B), collaterals (C), and boutons (D) per DL1 PN across developmental time. Statistics: one-way ANOVA followed by Kruskal-Wallis test. Statistical relationships are summarized using the Compact Display System (CDS): groups sharing any CDS letter are not significantly different, whereas groups with no letters in common differ significantly. See Figure 4, supplemental table for detailed results. Each point represents one cell.

12hr APF is the first developmental moment after metamorphosis when DL1 PNs send axonal projections into the calyx (Jefferis et al., 2004). At this timepoint, we observed single DL1 PN axons within the calyx that are highly decorated with filopodia. Some of these axonal filopodia contain small bouton-like swellings and extensions, while other axonal filopodia were naked. Strikingly, the number of axonal filopodia containing bouton-like swellings was already equivalent to the final adult number of collaterals, suggesting very early patterning of the final adult form (Figure 4).

DL1 PNs at 24hr APF did not have excess naked filopodia but rather an adult number of collaterals bearing immature boutons. These immature boutons were extensively decorated with filopodia, distinct from mature boutons which are compact and smooth. By 36hr APF, single DL1 PNs exhibit morphology that is already very similar to the final adult form. While ultimate collateral number was set early, bouton number increases until 36hr APF, coinciding with a qualitatively significant maturation of bouton structure and reduction in length of filopodia extensions found on boutons. By 48hr APF, DL1 morphology is qualitatively and quantitatively adult-like: bouton number has converged on final adult values, collateral number remains at adult levels, and bouton filopodia appear to have further retracted in length (Figure 4). After this time point, the DL1 axonal structure in the calyx grows—likely due to the continuing addition of Kenyon cells and inflation of the synaptic material in boutons—but does not change its form.

In sum, PN collateral number is fixed well before KC neurogenesis ends, implying that PNs rely on an early-emerging KC cue rather than final KC number to scale collateral numbers. Our data point to collateral growth and/or stabilization as the morphogenic step that implements bouton-number scaling.

### Projection neurons require Kenyon cells to grow and maintain nascent collaterals

In single DL1 PNs, the large excess of axonal filopodia present at 12hr APF is pruned to the adult number of collaterals by 24hr APF (Figure 4). This suggests that 12-24hr APF may represent a developmental window during which collateral production is sensitive to an unknown correlate of future KC population size. To test this, we used the broad PN driver *VT033006*-LexA, which labels most adPNs and lPNs, and manipulated KC number via mild HU treatment (Figure 5).

**Fig 5:**
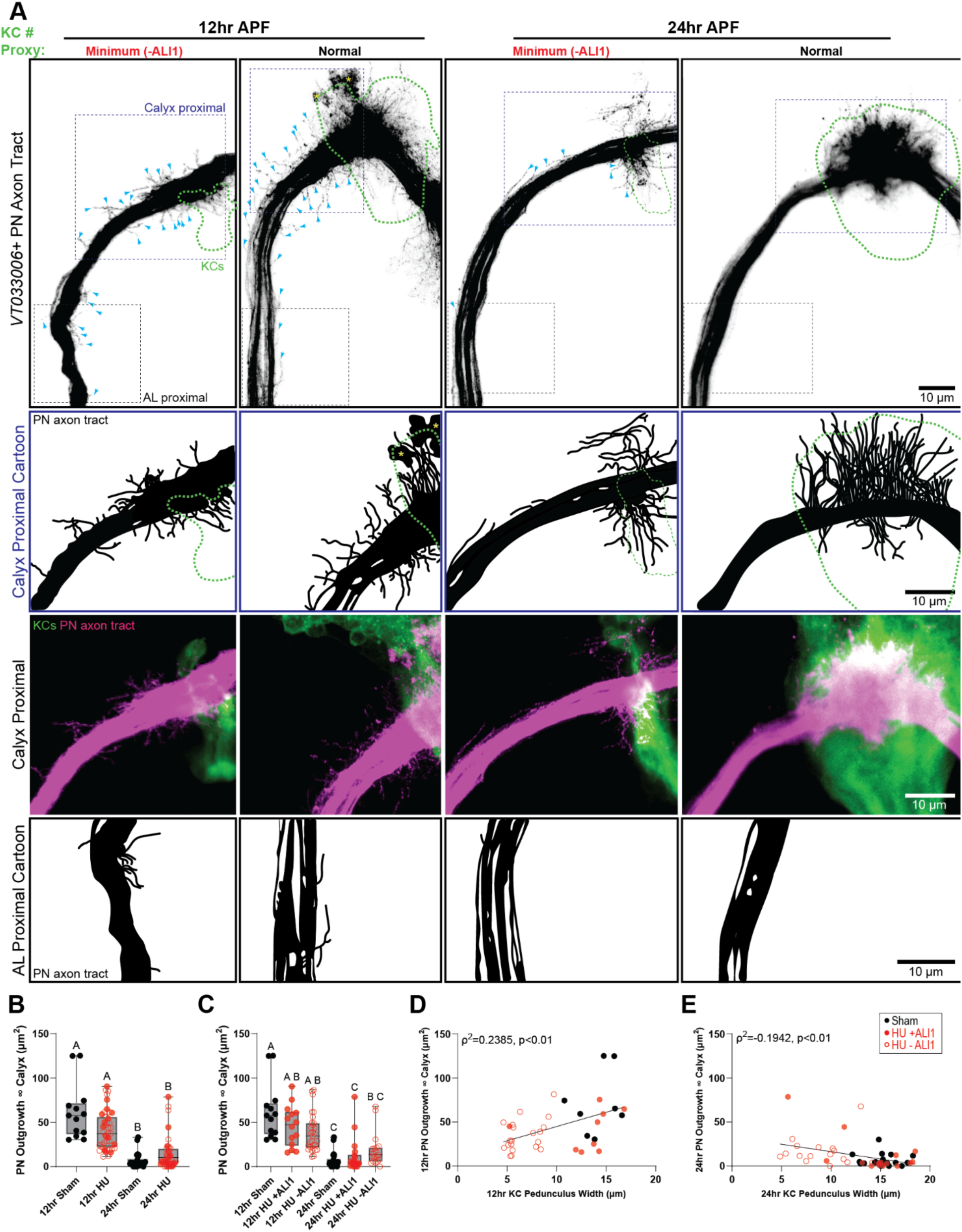
PN axonal filopodia transition from widespread exploration to calyx-restricted stabilization during early pupal development. **A**: (Row 1) Representative images of *VT033006*+ PN axon tracts at 12hr and 24hr APF. “Normal” samples are of the sham condition and have normal numbers of KCs. “Minimum −ALl1” samples lack all post-embryonically born KCs and ALl1. Green outline signifies the territory occupied by *OK107*+ KCs in each condition. Here and throughout: blue arrows: filopodial growth beyond the *OK107*+ KC territory; yellow asterisks: larval calyx debris. (Row 2) Cartoon representations of calyx proximal regions outlined in Row 1. Due to dense *VT033006* labeling, PN collaterals inside the calyx are shown as schematic sticks without bouton structures;.(Row 3) Raw fluorescence images of calyx proximal regions outlined with blue dashed boxes Row 1 and used to generate cartoons in Row 2 (*VT033006*+ PNs in magenta; *OK107*+ KCs in green). (Row 4) Cartoon representations of AL proximal regions outlined with black dashed boxes in Row 1. **B–C**: Box-and-whisker plots quantifying PN outgrowth beyond the calyx at 12hr and 24hr APF, comparing sham (black points) and HU (red points) conditions. (C) displays the same data as in (B), but now further separating HU brains into +ALl1 (solid red points) and −ALl1 (empty red points) categories. Statistics: one-way ANOVA followed by Kruskal–Wallis, with Compact Display System (CDS) lettering indicating significant group differences (see Figure 5 supplemental table for details). Each point is one calyx. **D–E**: Relationship between KC pedunculus width and PN outgrowth beyond the calyx at 12hr (D) and 24hr (E). Statistics: two-tailed Spearman’s correlation. Best-fit lines from simple linear regression. Each point is one calyx.

At 12hr APF in control brains, *VT033006* PNs display widespread filopodial outgrowth (“nascent collaterals”) along the axon tract, both within the calyx and well beyond it. Nascent collaterals within the calyx appear substantially longer than those outside the calyx, suggesting that KCs are permissive for collateral extension (Figure 5A). Consistent with this, nascent collateral outgrowth into the calyx scaled with calyx area at both 12hr and 24hr APF (Figure 5 Supplemental 1). Additionally, outgrowth beyond the calyx also appeared to positively scale with calyx area at 12hr but not 24hr APF, suggesting KCs may provide a secreted growth cue to PNs before 24hr APF (Figure 5A-E).

In hemispheres with complete loss of post-embryonic KCs (as well as the ALl1 lineage), nascent collaterals were still observed in the calyx at 12hr APF but were shortened relative to those of the sham-treated group (Figure 5A, Figure 5 Supplemental 1A). These shortened nascent collaterals often extended toward, and sometimes interacted with, the few remaining embryonically born KCs. Moreover, nascent collaterals located outside the calyx at 12hr APF were eliminated by 24hr APF in both control and HU-treated brains (Figure 5A, Figure 5 Supplemental 1A), suggesting that KCs provide signals required not only for nascent collateral growth but also for their local stabilization.

To determine whether collateral elimination in the absence of olfactory KCs persists into later development, we repeated the HU ablation experiments using a strong HU dose to reliably remove all KC NBs. Comparing ∼24hr APF (end of putative plasticity window) and ∼48hr APF (adult-like morphology) we again found that collateral sprouting into the calyx was severely disrupted at 24hr APF in KC-ablated hemispheres. Occasionally, a thin, elongated collateral bundle remained and appeared to interact with the embryonic KCs born before neuroblast ablation. By 48hr APF, the collateral depletion seen at 24hr APF had not recovered; instead, collaterals were completely absent from the calyx in KC-ablated hemispheres (Figure 6).

**Fig 6:**
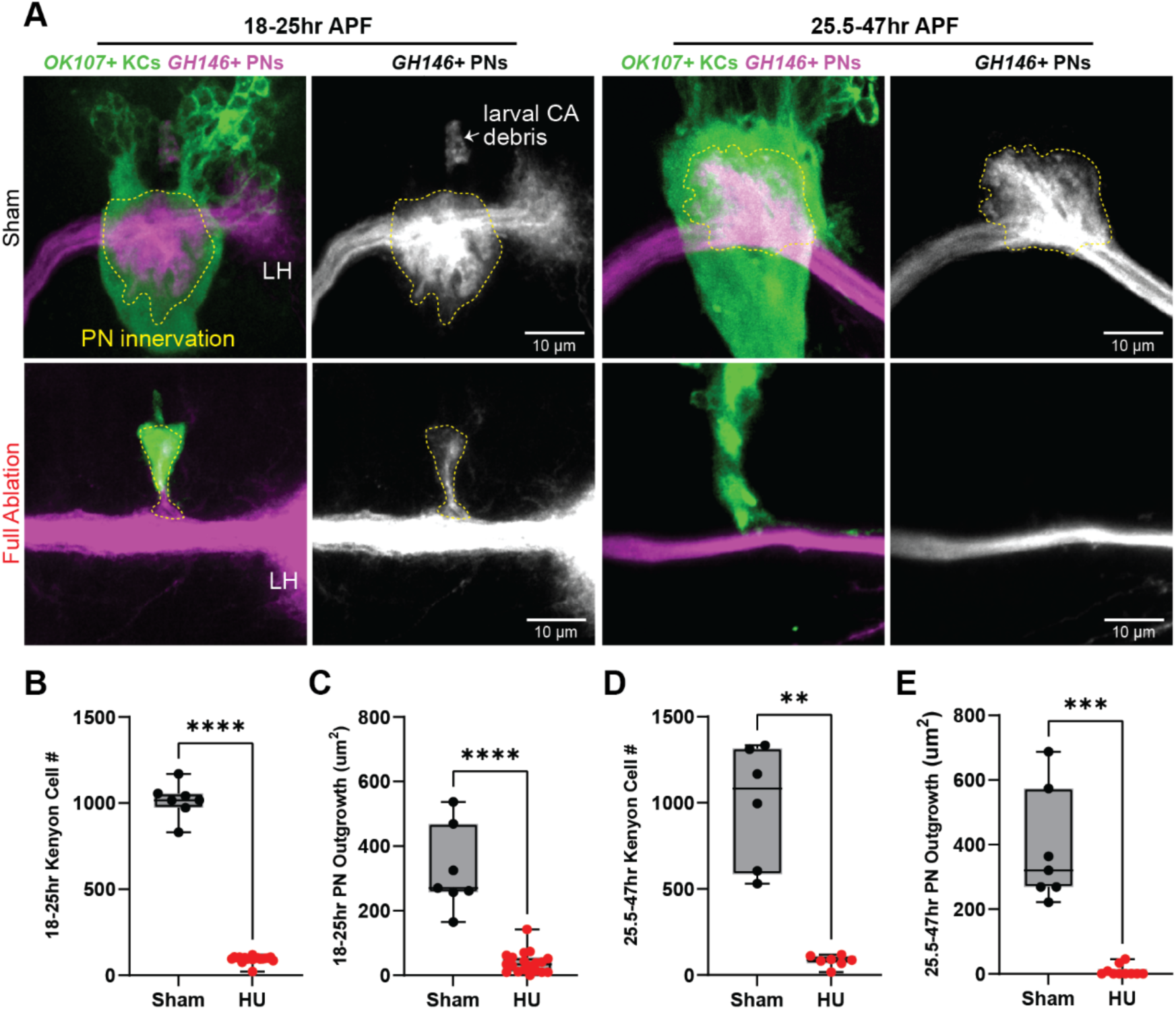
Post-embryonically born KCs are required for stabilization of PN axonal innervations into the calyx: Representative images of KCs (green) and PNs (magenta) at 18–25 hr APF (left) and 25.5–47 hr APF (right), shown for sham controls (top row) and strongly HU ablated samples lacking all 4 MBNBs (bottom row). Yellow dashed outlines mark PN innervation of the calyx. LH: lateral horn. CA: calyx. **B–C**: Box-and-whisker plots reporting KC number (B) or PN innervation of the calyx (C) at 18.5-25hr APF in sham and strong HU conditions. Each point represents a single calyx. **D–E**: Box-and-whisker plots reporting KC number (B) or PN innervation of the calyx (C) at 18.5-25hr APF in sham and strong HU conditions. Each point represents a single calyx. Statistics: two-tailed Mann–Whitney test; ****p < 0.0001, ***p<0.001, **p<0.01.

### Variation in Kenyon cell number drives projection neuron morphological growth

Our data thus far suggests a model in which PNs of all types grow and maintain collaterals in response to KC-derived cues. To explore if and how other dimensions of PN morphology change to accommodate variable numbers of KCs, we quantified a variety of morphometric parameters for VM4 and DL1 adPNs. We excluded VM6 adPNs as our labeling method for this PN subtype included other distracting cells in the calyx region.

PN type–specific bouton placement along the dorsal–ventral calyx axis is predicted by collateral length, so we first asked how collateral morphology scales with KC number. VM4 collateral lengths were unchanged across conditions and did not vary with KC population size, whereas DL1 collaterals were reduced in HU brains and scaled strongly with KC number. Higher-order branches disperse boutons beyond the main collateral. Lengths of higher-order branches were largely invariant in both PN types, though both VM4 and DL1 showed reduced branching below particular KC number thresholds. To capture total presynaptic growth, we combined collateral and branch lengths and found that total skeleton length scaled robustly with KC number in both PN types, reflecting increased production of modular bouton units (i.e., additional collaterals) as KC number rises. In contrast, the medial–lateral spacing of collaterals remained constant across KC conditions, indicating at least a degree of preserved spatial patterning of boutons. Bouton size was also largely invariant, though each PN subtype showed a weak positive relationship between bouton area and KC population size, while total bouton membrane per cell scaled strongly with KC number. The latter likely reflects the scaling of bouton number to KC population size (Figure 1 Supplemental 1-2; Figure 2 Supplemental 2-3).

Taken together, these analyses further support the conclusion that KC number shapes PN morphology primarily by regulating how collaterals are generated and maintained on a single PN in development. That spacing between bouton units, bouton number per collateral, bouton size, and overall relative number of collaterals on individual PNs are invariant to KC numbers suggests that at least some dimensions of morphological stereotypy are intrinsic to PNs.

### Developmental dynamics of projection neuron bouton filopodia suggest PN-PN interactions sculpt bouton morphologies

In addition to labeling a single cell born at the time of heat shock, tsMARCM also labels in a second color all sister cells born from the same clone after the moment of labeling induction. In inspecting these images, we noticed that boutons of both single DL1 PNs and their sister PNs were decorated with filopodia throughout development (Figure 7A). This prompted us to look qualitatively for whether bouton filopodia ever extended toward or contacted other boutons.

**Fig 7:**
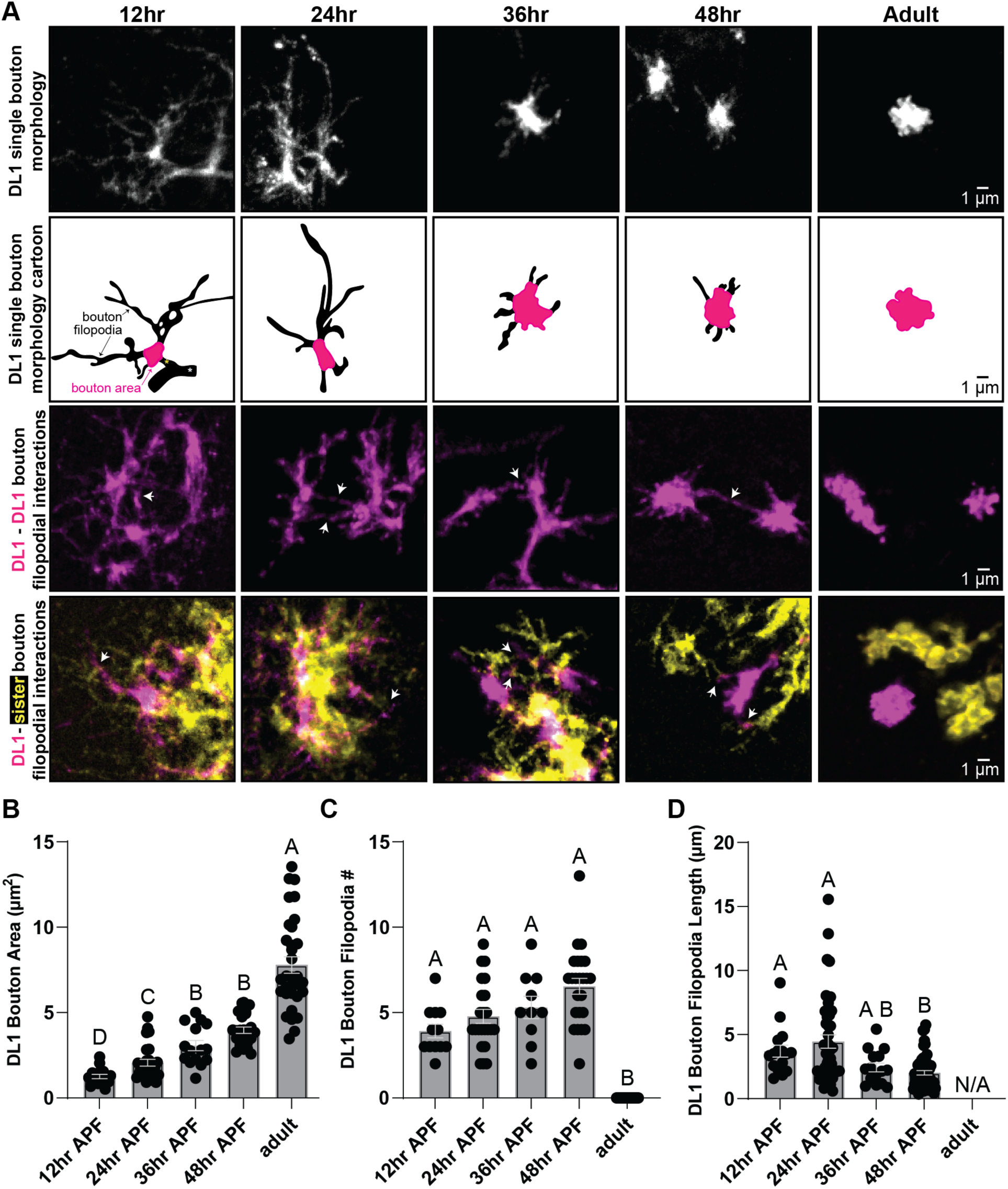
Transient interactions between bouton filopodia accompany bouton growth during maturation of DL1 adPN collateral-bouton assemblages. **A:** Top row: single boutons from developing DL1 PNs. Second row: schematic depictions of single boutons from top row. Magenta: representative bouton area at each developmental timepoint. Yellow asterisk marks the collateral; white asterisk marks the axon tract (12hr). Third row: white arrows indicate examples of interactions between neighboring DL1 bouton filopodia during development. Bottom row: white arrows indicate interactions between bouton filopodia of neighboring DL1 and twin-spot MARCM adPN clones across development. **B–D:** Box-and-whisker plots showing bouton area (B), filopodia number (C), and filopodia length (D) per DL1 PN over developmental time. Statistics: log-normal Brown–Forsythe and Welch ANOVA (B), or one-way ANOVA followed by Kruskal–Wallis test (C, D). Statistical relationships are summarized using the Compact Display System (CDS): groups sharing any CDS letter are not significantly different, whereas groups with no letters in common differ significantly. See Figure 5, supplemental table for detailed results. Each point represents one cell.

Across all developmental timepoints examined, bouton filopodia arising from DL1 collaterals extended toward and touched neighboring boutons or bouton filopodia. These interactions occurred both within the same DL1 PN and between DL1 boutons and boutons of sister PNs (Figure 7A). While filopodia number remained relatively stable, filopodia length underwent a marked transition: long exploratory filopodia at 12hr and 24hr APF shortened substantially by 36hr and 48hr APF and disappeared entirely in adults (Figure 7A,C,D). This coincided with a progressive increase in bouton area (i.e. a proxy for bouton volume), consistent with bouton maturation (Figure 7A,B).

In adulthood, boutons do not touch. Their separation could emerge through KC dendrites enwrapping and smoothing them, from PN-PN interactions, or both. Filipodial contact between boutons at early developmental timepoints, and the collapse of these filopodia coincident with expansion of bouton internal structure, suggests that repulsive interactions among PNs, or across boutons of the same PN, contributes to the emergence of bouton tiling. This would allow diffuse membranous structures to resolve into discrete units.

## Discussion

Ecologically unique selective pressures can reshape associative learning circuits by altering developmental programs, including those that determine how many principal neurons a circuit contains. However, the developmental mechanisms by which neurons detect and adjust to variation in their partner populations remain poorly understood. To probe these principles, we experimentally reduced KC number by HU ablation of KC NBs and examined how presynaptic PNs adjust their axonal morphogenesis in response. Building on our previous finding that adult PN bouton number is plastic and scales with experimentally increased or decreased KC number, we asked how these scaling mechanisms are implemented during development (Elkahlah et al., 2020).

Here, we find that flexible bouton production by PNs is driven by three developmental events. First, elongation of very early axonal filopodia (i.e. nascent collaterals) is sensitive to KC number. Second, only some nascent collaterals that sprout can mature and stabilize. And third, stabilized collaterals most often make just one bouton each, independent of PN identity or KC population size. Thus, PNs scale bouton number via the addition or elimination of modular collateral::bouton assemblages. This is both in response to KC population size and to achieve PN subtype-specific bouton counts. We additionally observed bouton-bouton interactions via bouton filopodia occurs within and across PN subtypes throughout development, which we posit supports resolving early membranous swellings into discrete, multi-synaptic boutons.

We observe PNs sprout nascent collaterals along the entire extent of the axon tract at 12hr APF, even in the complete absence of all four KC NBs. Furthermore, the degree of nascent collateral sprouting positively scales with KC number, with greater density and elongation of nascent collaterals occurring closer to the calyx core. This finding supports the idea that KCs secrete a permissive, diffusible cue that promotes nascent collateral outgrowth and forms a gradient extending beyond the calyx. Notably, embryonically born KCs on their own can also support limited sprouting of nascent collaterals, but not their stabilization. This suggests that additional signals, likely supplied by post-embryonically born KCs, are required to maintain mature collateral::bouton units.

Axon collateral formation is known to be a stochastic, multi-step process coordinated by mitochondrial positioning and respiration as well as cytoskeletal protein dynamics (Armijo-Weingart and Gallo, 2017). Previous work from our group demonstrated that PN bouton number scales not only with KC population size but also with the number of dendrites per individual KC (Ahmed et al., 2023), further indicating that bouton morphogenesis is sensitive to quantitative properties of the postsynaptic environment. Considering our findings that PNs become sensitive to KC-derived cues soon after pupation and establish largely adult-like collateral organization well before KC neurogenesis is complete, several, non-mutually exclusive mechanisms could underlie how PNs anticipate their final bouton complement. First, PNs may measure the total amount of KC-derived material present in the early calyx, such as (1) the number or extent of nascent KC dendrites, or (2) debris from the larval calyx, and use one or more of these signals to bias the probabilistic stabilization of axonal filopodia into mature collaterals. Second, PNs may be sensitive to the number of KC neuroblast clones contributing to the calyx at early pupal stages, using a correlate of clone number rather than absolute KC count as a proxy for the eventual size of the KC population. Third, PNs may respond preferentially to signals derived from specific KC subtypes whose presence or absence could disproportionately influence collateral stabilization. Determining whether one or multiple of these mechanisms occurs will require future experiments that independently manipulate KC clone number, KC number per clone, or KC subtype composition in advance of early pupal development (Elkahlah et al., 2020; Schlegel et al., 2024).

Individual collaterals most often have one bouton and almost never exceed three boutons. Collateral::bouton units define PN subtype bouton number, are robust to variable KC number, and scale to KC population size. These findings raise the possibility that a universal constraint shared by PN subtypes defines how many presynaptic sites any single collateral can support. This could be a cell-autonomous genetic growth program, or result from the progression of shared developmental events commonly constrained by time and space. The genome likely does not encode bouton number per collateral directly but instead executes a developmental algorithm which places strong constraints on the cytoskeletal patterning that can take place within a nascent collateral. We additionally provide evidence that PNs commit to an adult number of collateral::bouton units as early as 12hr APF, a time when KC neurogenesis is far from complete and the final number of dendritic claws cannot be known to PNs. One implication of this offset in developmental timing between PN morphogenesis and KC birth is that early bouton positioning might not require matching with specific KC partners. Instead, our observations are consistent with a model in which KCs later sample from a prepatterned array of bouton units, choosing boutons stochastically as their claws grow into the calyx. And, early tiling of boutons would ensure each KC claw can easily select just one bouton when the time for partnership selection arrives. If such a mechanism is employed it would naturally generate the sparse, varied PN::KC connectivity observed in the calyx, while still allowing bouton number to scale to the overall size of the KC population. Together, future experiments could test the following model: PNs specify calyx architecture (i.e. a scaffold of modular bouton units laid out early), and KCs specify the final wiring as they elaborate dendritic claws, thus allowing the circuit to remain robust to changes in KC population size.

## Supporting information

Supplemental table for Figure 3

Supplemental table for Figure 4

Supplemental table for Figure 5

Supplemental table for Figure 7

## Acknowledgements

We thank the Bloomington Drosophila Stock Center and Tzumin Lee for sharing fly strains; Ching-Po Yang for advice on twin-spot MARCM experimental set-up; and Clowney Lab members for providing input on the manuscript. V.M.P. was supported by K12-GM111725 and HHMI Hanna Gray Fellowship. E.M.T.-K. was supported by NIH T32 DC00011. E.J.C. is a McKnight Scholar, Pew Biomedical Scholar, and Rita Allen Milton Cassell Scholar. Funding was provided by NIDCD R01DC018032 (to E.J.C.).

## Methods

### Resources Table

| REAGENT or RESOURCE | SOURCE | IDENTIFIER |
| --- | --- | --- |
| Antibodies |  |  |
| anti-Acj6 (mouse) 1:50 | Developmental Studies Hybridoma Bank | Cat# anti-Acj6 |
| anti-ChAT (mouse monoclonal) 1:200 | Developmental Studies Hybridoma Bank | Cat# chat4b1; RRID: AB_528122 |
| anti-DsRed (rabbit polyclonal) 1:1000 | Clontech/Takara Bio | Cat# 632496; RRID: AB_10013483 |
| anti-GFP (chicken polyclonal) 1:5000 | Dawen Cai Lab | N/A |
| anti-GFP (rabbit polyclonal)<br>1:250 | Fisher | Cat# A11122 |
| anti-RFP (rat monoclonal)<br>1:250 | Chromotek | Cat# 5f8 |
| anti-mCherry (chicken)<br>1:1000 | Novus Biologicals | NBP2-25158-0.025ml |
| anti-CD8 (rat monoclonal)<br>1:100 | AbboMax | Cat# 603-062 |
| Alexa 488 anti-chicken (goat polyclonal) 1:1000 | Fisher | Cat# A32931; RRID: AB_2762843 |
| Alexa 488 anti-rabbit (goat polyclonal) 1:1000 | Fisher | Cat# A11008 |
| Alexa 568 anti-rabbit (goat polyclonal) 1:1000 | Fisher | Cat# A-11036; RRID: AB_10563566 |
| Alexa 568 anti-rat (goat polyclonal) 1:1000 | Fisher | Cat# A11077 |
| Alexa 647 anti-mouse (goat polyclonal) 1:1000 | Fisher | Cat# A-21236; RRID: AB_2535805 |
| Chemicals, peptides, and recombinant proteins |  |  |
| Hydroxyurea | Sigma | H8627 |
| Paraformaldehyde | EMS | 15710 |
| DAPI (used in stainings though not shown in figures) | Sigma | D9542 |
| 10X PBS Gibco | Fisher | 70-011-044 |
| Normal Goat Serum | MP biomedicals | 8642921 |
| Triton | Sigma | X100 |
| Sodium chloride NaCl | Sigma | S7653 |
| Potassium chloride KCl | Fisher | P217-500 |
| Calcium chloride<br>CaCl<sub>2</sub>*2H<sub>2</sub>O | Fisher | C70-500 |
| Magnesium chloride<br>MgCl<sub>2</sub>*6H<sub>2</sub>O | Sigma | M2670 |
| HEPES sodium salt | Sigma | H7006 |
| Trehalose | Sigma | T9531 |
| Sucrose | Sigma | 84097 |
| Sodium bicarbonate<br>NaHCO<sub>3</sub> | Fisher | S233-500 |
| Sodium phosphate<br>NaH<sub>2</sub>PO<sub>4</sub>*H<sub>2</sub>O | Fisher | S369-500 |
| 1X Dulbecco's PBS | Corning | 21-031-CV |
| Other |  |  |
| Fly food | Lab Express, Ann Arbor MI | “B” Food and “J Food” |
| Binder reinforcement stickers<br>(for mounting brains) | Office Depot | Avery 5722 |
| Strainer cap | Amazon | ASIN: B0D7LL9F4R |
| Software and algorithms |  |  |
| R | R Foundation for Statistical Computing | www.R-project.org |
| R package dbscan | [doi:10.18637/jss.v091.i01](https://doi.org/10.18637/jss.v091.i01) |  |
| Imaris | Oxford Instruments | RRID: SCR_007370<br>https://imaris.oxinst.com/packages |

### Fly husbandry

All fly lines used in this study are listed in the key resources table. Flies were maintained at 25°C on a 12:12 light-dark cycle with humidity provided by a beaker of water in the incubator. Fly stocks were maintained on Bloomington food (“B” recipe, Lab Express, Ann Arbor, MI) sprinkled with yeast. Larval density in development can impact KC population sizes observed in adult brains (Heisenberg et al., 1995; Lin et al., 2013). Thus, for experiments conducted on B food, the number of female and male parents were kept consistent across crosses and flipped consistently every 2 to 3 days. For tsMARCM experiments, larvae were collected in 2-hour windows immediately after hatching on grape plates sprinkled with yeast. Collected larvae were transferred in groups of 80 to Janelia food vials (“J” recipe, Lab Express, Ann Arbor) with a layer of pre-mushed food on top.

For pupal developmental staging, pupae were transferred using a wet paintbrush to new vials at the 0hr APF “white pupae” stage. Single vials were populated with newly pupated animals for windows of just 2 hours, over the course of up to 24 hours for any given experiment. Pupa were dissected at the required times defined by these 2-hour collection windows (i.e. if pupa from a vial collected at 0hr APF between 8am-10am were subsequently dissected two days later between 8am-10am, dissected brains would be given the developmental stage label of “48hr APF” – thus, “48hr APF” data actually captures stages 46-50hr APF).

### Fly genotypes by figure

| Figure | Genotype |
| --- | --- |
| Figure 1, Figure 1 Supplemental 1-2, Figure 3A-C, Figure 3 Supplemental 1G-I | $\frac{hsflp}{Y \text{ or } +}; \frac{UAS-rCD2.RFP,UAS-GFPi,FRT40A,GH146-gal4}{UAS-mCD8-GFP,UAS-rCD2i,FRT40A}; +$ |
| Figure 4, Figure 7 | $\frac{hsflp}{Y \text{ or } +}; \frac{UAS-rCD2.RFP,UAS-GFPi,FRT40A,GH146-gal4}{UAS-mCD8-GFP,UAS-rCD2i,FRT40A}; +$<br>Or<br>$\frac{hsflp}{Y \text{ or } +}; +; \frac{FRTG13,GH146-gal4,UAS-rCD2.RFP,UAS-GFPi}{FRTG13,GH146-gal4,UAS-mCD8-GFP,UAS-rCD2i}; +$ |
| Figure 2C-F, Figure 2 Supplemental 2-3, Figure 3A-C, Figure 3 Supplemental 1A-C<br>These data are re-analyzed from Elkahlah et al., 2020. | $;\frac{MB247dsRed}{CyO};\frac{42D01-Gal4,UAS-C3PA}{42D01-Gal4,UAS-C3PA};$ |
| Figure 2H-K, Figure 2 Supplemental 1, Figure 3A-C, Figure 3 Supplemental 1D-F. These data are re-analyzed from Elkahlah et al., 2020. | $;\frac{MB247dsRed}{CyO};\frac{71D09-Gal4,UAS-mCD8-GFP}{71D09-Gal4,UAS-mCD8-GFP};$ |
| Figure 5, Figure 5 Supplemental 1 | $;\frac{VT033006-LexA,LexAop-Tomato}{+};\frac{UAS-mCD8-GFP}{+};$<br>$\frac{OK107-Gal4}{+}$ |
| Figure 6 | $;\frac{GH146-QF,QUAS-mtdTomato-3xHA}{+};\frac{UAS-mCD8-GFP}{+};$<br>$\frac{OK107-Gal4}{+}$ |

### Fly alleles

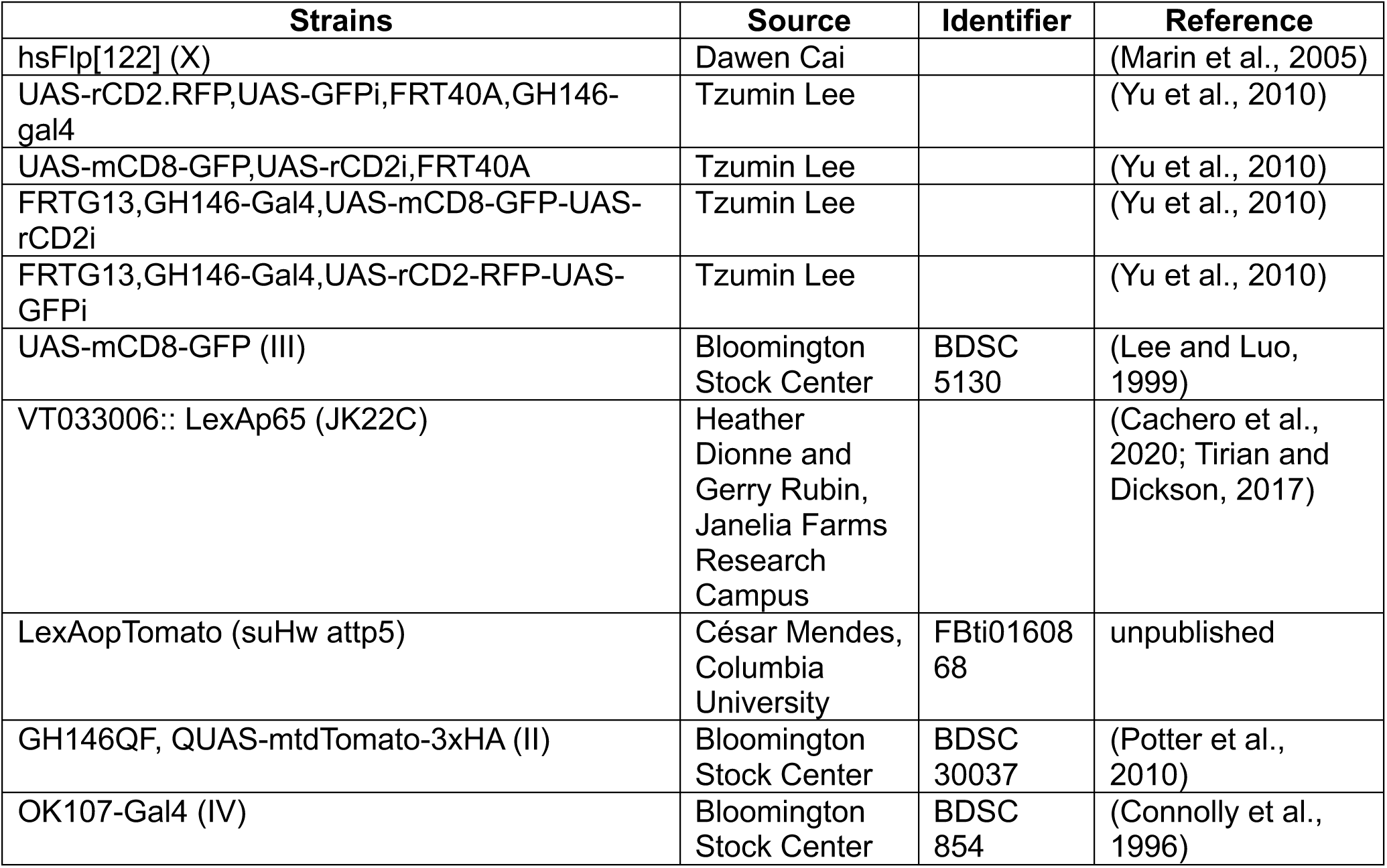

| Strains | Source | Identifier | Reference |
| --- | --- | --- | --- |
| hsFlp[122] (X) | Dawen Cai |  | (Marin et al., 2005) |
| UAS-rCD2.RFP,UAS-GFPi,FRT40A,GH146-gal4 | Tzumin Lee |  | (Yu et al., 2010) |
| UAS-mCD8-GFP,UAS-rCD2i,FRT40A | Tzumin Lee |  | (Yu et al., 2010) |
| FRTG13,GH146-Gal4,UAS-mCD8-GFP-UAS-rCD2i | Tzumin Lee |  | (Yu et al., 2010) |
| FRTG13,GH146-Gal4,UAS-rCD2-RFP-UAS-GFPi | Tzumin Lee |  | (Yu et al., 2010) |
| UAS-mCD8-GFP (III) | Bloomington Stock Center | BDSC 5130 | (Lee and Luo, 1999) |
| VT033006:: LexAp65 (JK22C) | Heather Dionne and Gerry Rubin, Janelia Farms Research Campus |  | (Cachero et al., 2020; Tirian and Dickson, 2017) |
| LexAopTomato (suHw attp5) | César Mendes, Columbia University | FBti0160868 | unpublished |
| GH146QF, QUAS-mtdTomato-3xHA (II) | Bloomington Stock Center | BDSC 30037 | (Potter et al., 2010) |
| OK107-Gal4 (IV) | Bloomington Stock Center | BDSC 854 | (Connolly et al., 1996) |

### Twin-spot MARCM

For twin-spot MARCM visualization of single cells throughout development and in the adult, larval and pupal development were synchronized as described in “Fly Husbandry”. To induce PN clones, larva were subjected to heatshock at 37°C for 30 minutes and then returned to 25°C until dissection.

DL1 PNs arise from the ALad1 lineage, and are the only adPN type generated during the first 24 hours after larval hatching (Jefferis et al., 2004; Wu and Luo, 2006). We took advantage of this developmental feature of DL1 PNs and used tsMARCM to visualize the morphogenesis of individual DL1 PNs born between 18.5 and 21 hours ALH. We verified previous reports that the only adPNs generated in this window are DL1 PNs: 100% of adult brains with adPN clones contained single PNs projecting dendritic arbors to the DL1 glomerulus. In developing brains, we distinguished lPN and adPN clones from each other by the absence or presence of anti-Acj6 nuclear labeling, respectively (Komiyama et al., 2003). Previous reports demonstrate that dendrites of developing PNs project to the approximate territory of final adult glomeruli (Lyu et al., 2025). We used this as the basis for interpreting the following finding: single PNs observed within Acj6+ clones projected their dendrites to AL territories similar to the adult location of the DL1 glomerulus at each pupal time point analyzed. Taken together, we had high confidence that the single PNs observed throughout development were consistently of the DL1 identity.

For tsMARCM experiments used to visualize developing DL1 PNs, FRT40A and FRTG13 alleles produced identical results (data not shown) and thus data were combined in the work presented here.

### Hydroxyurea ablation of Kenyon cells

Protocol was adapted from (Sweeney et al., 2012). Genesis of data in Figure 2 is described in Elkahlah et al., 2020. Experiments in Figure 6 were conducted using the workflow described in Elkahlah et al., 2020, except that we treated animals with 10mg/mL hydroxyurea for 4 hours to ablate all KC neuroblasts.

For Figures 1 and 5, we set up large populations of flies in cages two days prior to ablation and affixed a homemade 35 mm grape juice agar plate containing yeast paste as a lid to the cage. We allowed flies to acclimate to the cage and yeasted grape plate for 2 days, replacing the yeasted grape plate each day. On day 3, we again replaced the yeasted grape plate with a fresh one and began embryo collection. On the morning of the ablation, we removed the plate from the cage and allowed larvae to hatch over a maximum of 6 hours. Larvae were transferred off the plate via pipetting in sucrose solution to a strainer cap submerged in a dish containing sucrose solution. Larvae were kept at a density of 80 larvae per submerged strainer cap. Larvae were then transferred using the strainer cap either to a dish containing hydroxyurea mixed in yeast:sucrose solution, or to a dish with sham yeast:sucrose mixture without HU. For both Figures 1 and 5, we used 12.5 mg/mL for 45-60 minutes to obtain variable KC neuroblast ablation across animals.

Following ablation or sham conditions, strainer caps with larvae were transferred to dishes containing sucrose solution for a quick rinse. Quick rinses were repeated twice more in separate dishes with fresh sucrose solution. Strainer caps with larvae were then transferred to a final dish containing sucrose solution, and larvae were then pipetted from the strainer cap and moved to coffee filter “tents” in order to separate larvae from the sucrose solution. Coffee filter “tents” containing larvae were placed in a vial of J food (“J” recipe, Lab Express, Ann Arbor) containing a layer of pre-mushed food at the surface.

We note that only the ALl1 NB in addition to KC NBs are sensitive to HU ablation within the first few hours after larval hatching, as we and others have previously documented (Elkahlah et al., 2020; Lovick and Hartenstein, 2015). Because loss of other PNs increases bouton production by spared PNs, while loss of KCs decreases bouton production, differences in inter-PN competition across samples may weaken some of the trends we describe here (Elkahlah et al., 2020; Thornton-Kolbe et al., 2025).

For our re-analysis of previously published *R42D01*-Gal4 data (Elkahlah et al., 2020), we selected only samples in which VM4 was labeled, using lateral horn innervation patterns to distinguish photoactivated VM4 PNs from VM3 PNs, as glomeruli from these PNs are easily distinguishable when single cells are photoactivated.

### Brain dissection, immunostaining, and confocal imaging

For immunostaining, brains were dissected in external saline (108 mM NaCl, 5 mM KCl, 2 mM CaCl2, 8.2 mM MgCl2, 4 mM NaHCO3, 1 mM NaH2PO4, 5 mM trehalose, 10 mM sucrose, 5 mM HEPES pH7.5, osmolarity adjusted to 265 mOsm) for up to 1 hour before being fixed. All steps were performed in 1.75 mL tubes or in cell strainer cap “baskets” in a 24-well plate. Brains were fixed for 25 minutes at room temperature in 4% PFA in 1X PBS and were subsequently washed 3x’s, 10 minutes/wash, in 1X PBST (1X PBS supplemented with 0.1% triton-x-100) on a shaker at room temperature, blocked 1 hour in 5% Normal Goat Serum in PBST, and then incubated for at least two overnights in primary antibody solution diluted in 5% Normal Goat Serum in PBST. Primary antibody was washed 3x’s, 10 minutes/wash, in PBST on a shaker at room temperature, then brains were incubated in secondary antibodies diluted 5% Normal Goat Serum in PBST for at least two overnights. DAPI (1 microgram/mL) was included in secondary antibody mixes. Antibodies and concentrations can be found in the resources table.

Brains were mounted in either a homemade mounting medium (1x PBS, 90% glycerol supplemented with propyl gallate) or with SlowFade™ Glass Soft-set Antifade media (Fisher) within binder reinforcement stickers sandwiched between two coverslips. The coverslip sandwiches were taped to slides, allowing us to perform confocal imaging on one side of the brain and then flip over the sandwich to allow a clear view of the other side of the brain. Samples were stored at room temperature in the dark prior to imaging.

Scanning confocal stacks were collected along the anterior-posterior axis on a Zeiss LSM 880 or Leica SP8 with 0.5 micrometer spacing in Z and 70nm axial pixel size, using a 63x 1.4 NA objective (Zeiss) or ∼150nm axial pixel size using a 40x 1.3 NA objective (Leica).

### Image analysis

<u>Analysis considerations:</u> We used mixed-sex populations. Sex differences in neuronal anatomy within the fly are well-documented. However, these have not been observed in the neuronal populations we study here. Any brains that appeared damaged from dissections, or those with the mushroom body region obscured due to insufficient tracheal removal, were not included in the analysis.

<u>Quantifying Kenyon cell number or its proxy:</u> When quantifying KC number directly, KCs were labeled using a cell type–specific GAL4, and KC somata were identified as DAPI+ cell bodies connected to the mushroom body. Labeled KC somata were counted using the FIJI multipoint tool in every third optical section (1 μm spacing between slices), using DAPI to distinguish individual cells.

To obtain a proxy for KC number, we used two approaches depending on the experimental context.

1. tsMARCM experiments: In experiments where adult PNs were labeled using tsMARCM, genetic labeling of KCs was not possible. For hemispheres in which only ALad1 clones were labeled (i.e., without simultaneous ALl1 labeling), we quantified the calyx area occupied by boutons from the labeled adPNs. We identified the Z-plane with the largest ALad1 clone–associated calyx cross-sectional area, manually outlined this region in FIJI, and measured its area using the “Measure” function.
2. All other experiments: When KC labeling was available via a cell type–specific GAL4, we estimated KC number by measuring KC-labeled calyx size. For each hemisphere, we identified the Z-plane with the largest KC-labeled calyx cross-sectional area, outlined the calyx in FIJI, and quantified the area using the “Measure” function. In developmental datasets, we also used pedunculus width as a proxy for KC population size. Pedunculus width was measured within 20 µm of the calyx base, in Z-planes where the pedunculus exited the calyx in a near-parallel trajectory (a condition met in the majority of developmental images).

<u>Quantification of projection neuron bouton and collateral numbers:</u> Boutons, collaterals, and bouton number per collateral were quantified only in datasets where single PNs were individually labeled. In most experiments (adult and developmental), boutons were identified morphologically as discrete membranous swellings along PN collaterals; in Figure 2H–K, bouton identity was confirmed using both morphology and ChAT immunostaining. Single PN collaterals were defined as branches extending from the main PN axon tract within the mushroom body (MB) calyx and that bore one or more boutons. Boutons located directly on the primary axon tract (i.e., not attached to an identifiable axonal branch) were also scored as collaterals but of zero length.

In developing PNs, some timepoints exhibited axonal branches that lacked boutons; these bouton-negative branches were categorized and quantified as axonal filopodia. Axonal branches bearing bouton-like structures were classified as both collaterals and axonal filopodia.

Quantification of projection neuron morphological features from light microscopy data:

**Collateral length:** adult collateral length was measured from single PNs. Collaterals were defined as bouton-bearing side branches emerging from the main PN axon shaft within the mushroom body (MB) calyx. For each collateral, the path from its point of emergence on the primary axon to its distal most point was traced using either Imaris or FIJI.

**Higher-order branch length:** adult higher-order branches were defined as bouton-bearing extensions emerging from collaterals or other higher order branches. Higher order branch lengths were quantified by tracing each branch from its emergence point to the branch tip using Imaris.

**Total skeleton length:** adult total skeleton length per PN was calculated as the sum of all collateral lengths and higher-order branch lengths for the cell. The primary axon shaft was not included in this measurement.

**Single bouton area:** in both adult and developing single PNs, single boutons were identified morphologically as discrete membrane swellings along collaterals. For each bouton, the Z-plane containing the largest cross-sectional area was selected, the bouton was outlined manually in FIJI, and area was measured using the “Measure” function.

**Total bouton area:** adult total bouton area per PN was quantified by summing cross-sectional bouton areas across all boutons belonging to a single PN.

Spacing between collaterals: spacing between collaterals in adult single PNs was measured along the main PN axon. For each adjacent pair of collaterals, the distance between their points of emergence on the primary axon was measured using Imaris.

Bouton filopodia number and length: bouton filopodia were identified by morphology as thin projections emanating from developing boutons. At each developmental timepoint, the number of filopodia extending from each bouton was counted manually using FIJI. Filopodia arising from collateral shafts rather than boutons were excluded from this metric. Filopodia length was measured from their point of emergence on the bouton surface to the distal tip of the filopodia. Measurements were made using FIJI with the segmented line tool.

<u>Quantification of projection neuron developmental outgrowth</u>: PN outgrowth was quantified by measuring the extent of VT033006+ or GH146+ PN axonal innervation of either the mushroom body (MB) calyx or beyond the calyx along the axon tract. To measure PN outgrowth inside the calyx, we manually outlined the PN-labeled axonal innervation of the calyx (defined by OK107+ KC dendrites) using FIJI and quantified the area using the “Measure” function. To measure PN outgrowth beyond the calyx, we identified PN filopodia that branched off of the axon tract between the calyx and antennal lobe. Using FIJI, we manually outlined and measured the area of each individual filopodia (or filopodial clusters if many were bundled together) found along the axon tract

### Statistical considerations

Brains were prepared for imaging in batches of 5–30. Genotypes or conditions being compared with one another were always prepared for staining together and imaged interspersed with one another to equalize batch effects. We excluded from analysis samples with overt physical damage to the cells or structures being measured.

Statistical tests and corresponding p-value significance thresholds are reported in each figure legend or directly on figures. For comparisons between two groups, two-tailed Mann–Whitney tests were used when data did not meet Gaussian distribution criteria; otherwise, two-tailed t-tests were applied. For correlation analyses, two-tailed Spearman’s rank correlations were used for non-Gaussian datasets, while Pearson correlations and simple linear regression were applied when appropriate. Figure legends specify which test was used for each dataset.

### Analysis of Electron Microscopy Connectomes

Boutons and collaterals were annotated for uniglomerular projection neurons in both the hemibrain connectome (Scheffer et al., 2020) and FAFB connectome (Dorkenwald et al., 2024; Schlegel et al., 2024; Zheng et al., 2018). Annotations for Hemibrain data were done in Thornton-Kolbe et al., 2025. In short, 24,791 synapses from 105 uniglomerular olfactory PNs were clustered into 538 boutons with a minimum of 3 synapses using k-means clustering and manual proofreading. 15 PNs from this data set had synapses with olfactory KCs just outside the defined calyx ROI. Thus, 361 synapses between PNs and KCs were added and clustered into 15 new boutons for the analyses presented here. Location of collateral branch points from the main axon were also annotated in Thornton-Kolbe et al., 2025. In short, collateral branch points were branch points on the EM reconstructed skeleton found along the “spine” (the longest continuous track) that gave rise to collaterals bearing synapses with KCs. 273 collateral branch points were annotated in Thornton-Kolbe et al., 2025 and 15 were added for these analyses.

Neuron skeletons, synapse coordinates, and cell types were downloaded from Codex (Dorkenwald et al., 2024; Schlegel et al., 2024; Zheng et al., 2018). 204 uniglomerular, olfactory PNs with synapses in the “CA ROI” were annotated. Coordinates of synapses between these 204 PNs and any cell in the “Kenyon Cell” class were used. We downloaded this data prior to July 2025 and thus used the original version of the data in which synapses were detected using a method described by Buhmann and colleagues and refined by Heinrich and colleagues (Buhmann et al., 2021; Heinrich et al., 2018). Synapses were clustered into boutons based on their spatial location using a density-based clustering method with the dbscan R package (Hahsler et al., 2019). Lone synapses (those not clustered) joined boutons within 1500nm, all others were not further analyzed. Boutons within 1500nm of one another that shared all or all but one KC partner were merged into single boutons. Abnormally small boutons, those with less than 5 KC partners were removed from further analysis. As in the hemibrain, collateral branch points were identified as those which branched from the “spine” segment and bore synapses with KCs. Boutons were assigned to collaterals by proximity of their coordinate centers to points on the collateral skeleton. For boutons located directly on the spine, collateral branch points were assigned as the closest point along the spine to the coordinate center of the bouton. In sum, for the two hemispheres of the FAFB, 326,940 synapses were clustered into 892 boutons on 587 collaterals. Each olfactory PN type was annotated with its type/ glomerular target in the AL (Dorkenwald et al., 2024). We further added neuroblast origin, developmental stage of birth, and birth order information from Lin et al., 2012 and Yu et al., 2010.

When generating scatter plots of bouton number versus collateral number from EM datasets, PNs whose absolute residual from the “All PNs” regression line fell within the top 5% of residuals and thus significantly deviated from expectation were called outliers.

---

**Fig 1, Supplemental 1:**
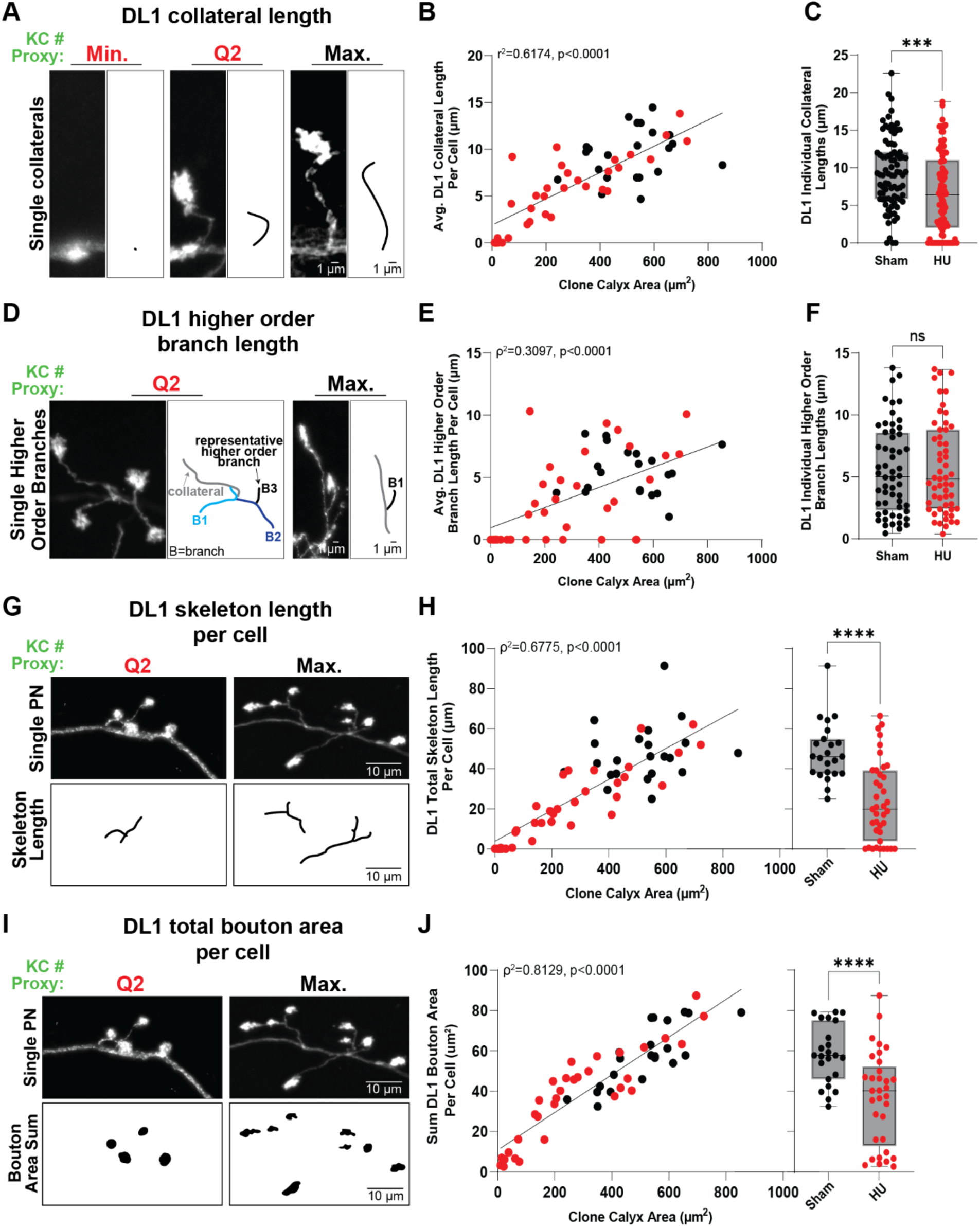
Multiple DL1 morphological features scale positively with KC population size. **A–C:** (A) Representative adult DL1 PNs over varying KC population sizes induced by HU ablation (red) or sham control (black). Cartoons depict traced collaterals (B) Scatter plots showing the relationship between clone calyx area and average DL1 collateral length per PN. Each point represents the average collateral length for one cell. Statistics: simple linear regression. (C) Box-and-whisker plots of DL1 collateral length across conditions. Each point represents a single collateral. Statistics: two-tailed Mann–Whitney test; ***p < 0.001. **D-F:** (D) Representative higher-order branches from adult DL1 PNs. Cartoons depict traced branches. (E) Scatter plots of clone calyx area versus average higher-order branch length per DL1 PN. Each point represents average higher order branch length for one cell. Statistics: two-tailed Spearman’s correlation. Best-fit lines from simple linear regression. (F) Box-and-whisker plots of higher-order branch length across conditions. Each point represents a single higher order branch. Statistics: two-tailed Mann–Whitney test; ns = 0.9688. **G-H:** (G) Representative adult DL1 PN skeleton lengths (i.e. the sum of all collateral and higher-order branch lengths per cell). Cartoons depict traced skeletons. (H) (Left) Scatter plot of clone calyx area versus total DL1 skeleton length. Each point represents a single cell. Statistics: two-tailed Spearman’s correlation. Best-fit lines from simple linear regression. (Right) Box-and-whisker plot of total skeleton length across conditions. Each point represents a single cell. Statistics: two-tailed unpaired t-test; \*\*\*\**p* < 0.0001. **I-J:** (I) Representative adult DL1 PNs showing bouton area sums per cell across KC population sizes. Cartoons depict traced bouton areas. (J) (Left) Scatter plot showing clone calyx area versus bouton area sum per DL1 PN. Each point represents a single cell. Statistics: two-tailed Spearman’s correlation. Best-fit lines from simple linear regression. (Right) Box-and-whisker plot of total bouton area per DL1 PN. Each point represents a single cell. Statistics: two-tailed unpaired t-test with Welch’s correction; \*\*\*\**p* < 0.0001. Min.: Minimum. Q2: Quartile 2. Max.: Maximum.

**Fig 1, Supplemental 2:**
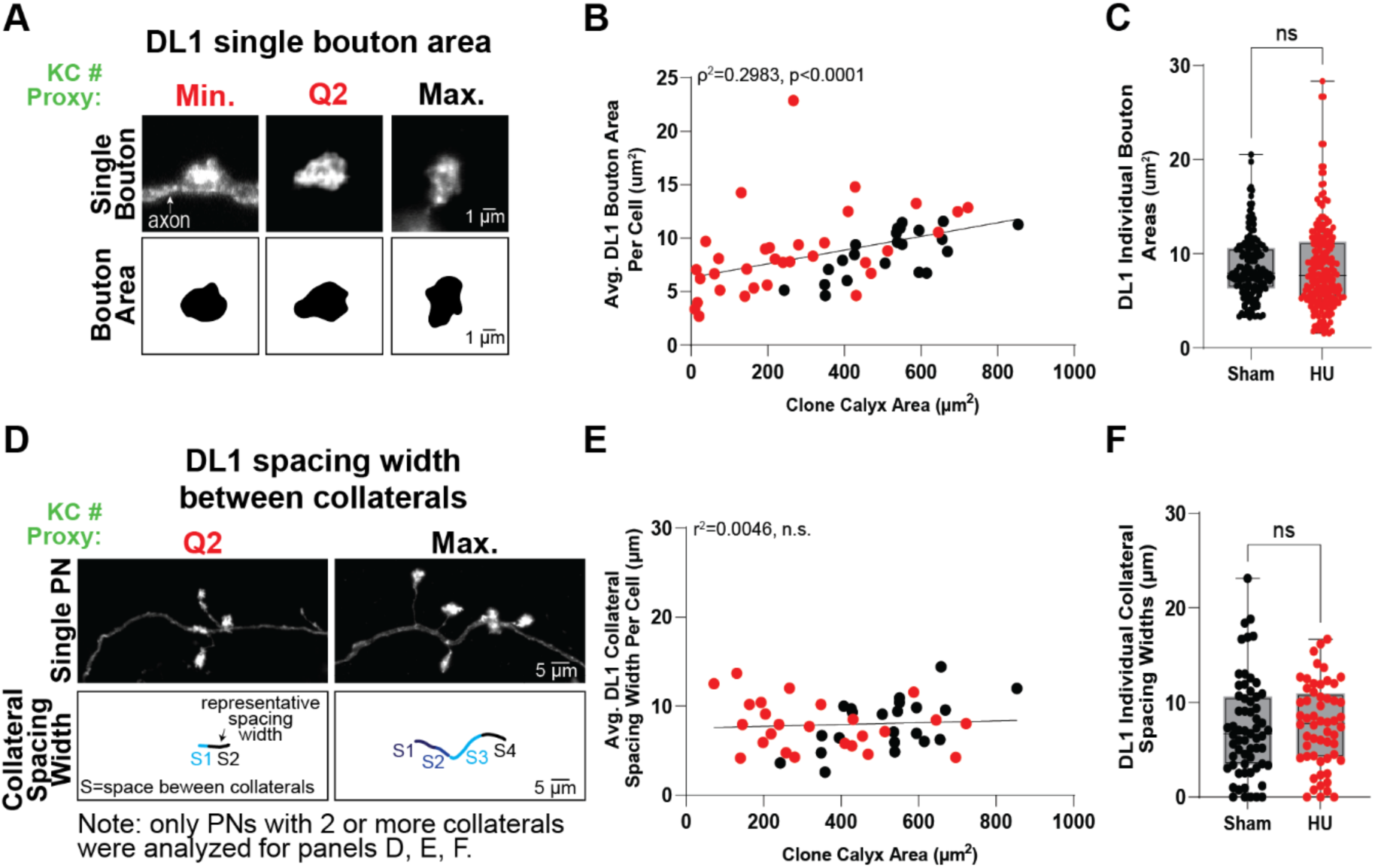
DL1 bouton size and collateral spacing show limited sensitivity to KC number. **A–C:** (A) Representative individual bouton areas for adult DL1 PNs across varying KC population sizes induced via HU ablation (red text), or sham control (black). Cartoons depict traced bouton areas. (B) Scatter plot showing the relationship between KC population size proxy and average bouton area for DL1 PNs. Each data point is the average bouton area for a single cell. Statistics: two-tailed Spearman’s correlation; best-fit lines from simple linear regression. (C) Box-and-whisker plot describing measures of bouton area for DL1 PNs across control (black dots) and HU (red dots) conditions. Each data point is a single bouton. Statistics: two-tailed Mann-Whitney test. n.s. p-value: 0.3787. **D-F:** (D) Relationship between KC population size and spacing width between individual collaterals along DL1 PN axons across HU (red) and control (black) conditions. Cartoons depict traced collateral spacing. (E) Scatter plot showing the relationship between KC population size proxy and average collateral spacing width per PN. Each point represents average space between collaterals for one cell. Statistics: simple linear regression. (F) Box-and-whisker plot of individual collateral spacing width for DL1 PNs across control (black) and HU (red) conditions. Each point represents one spacing measurement. Statistics: two-tailed Mann–Whitney test; n.s. p-value: 0.4674.

**Fig 2, Supplemental 1:**
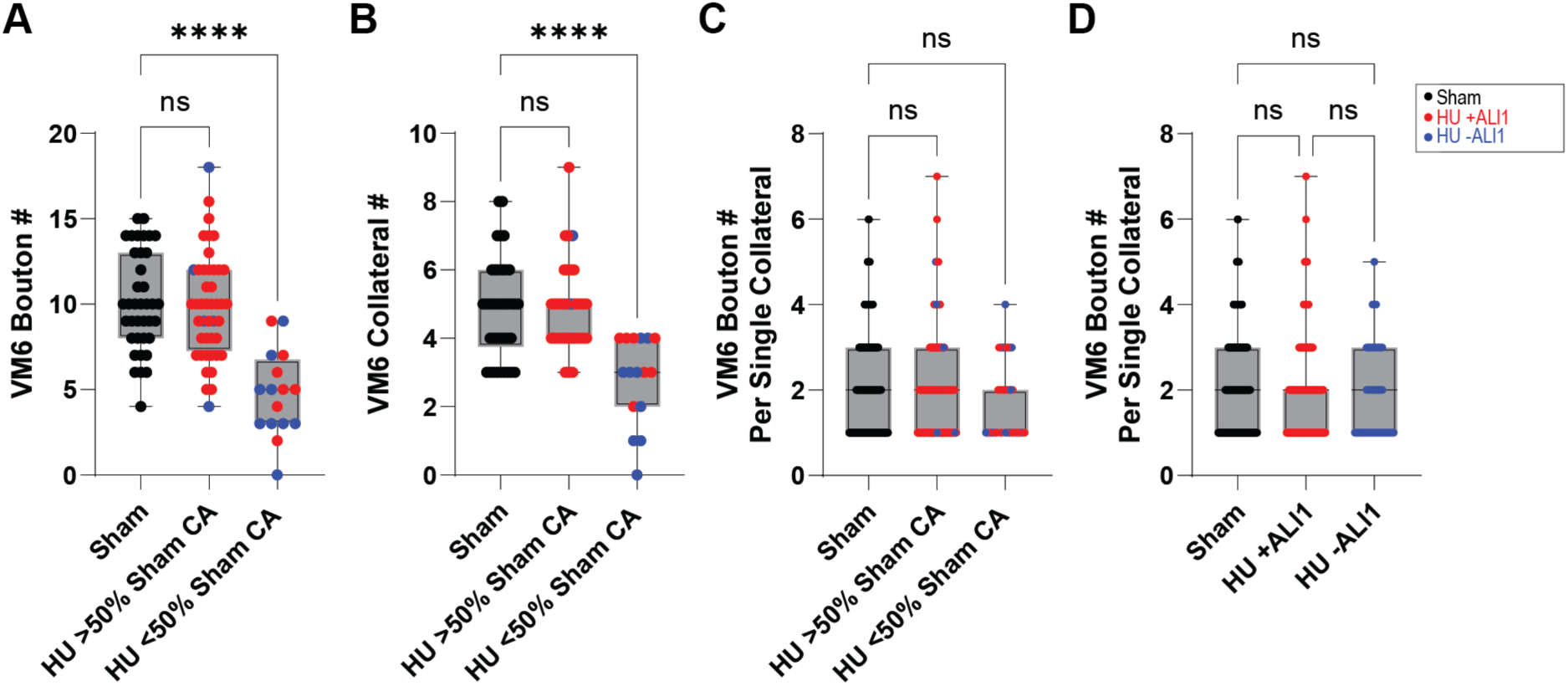
Smaller KC population sizes drive changes to bouton and collateral numbers in VM6 adPNs. **A-C:** Box-and-whisker plots describing measures of bouton number (A), collateral number (B), or bouton number per single collateral (C) for single *71D09*+ VM6 PNs across sham (black dots) and HU (red dots: +ALl1 blue dots: −ALl1) conditions, with HU data split by datapoints >50% sham average calyx area (CA), or <50% sham average CA. **D:** Box-and-whisker plot displaying the same data as in (C), but now HU data is split by datapoints with the ALl1 lineage either present (+ALl1) or absent (-ALl1). Each data point is a single collateral. Statistics: one-way ANOVA followed by Kruskal-Wallis test. \*\*\*\**p* < 0.0001. NS *p*-values: >0.9999 (A), >0.9999 (B), >0.9999 sham vs. HU >50%, or 0.0687 sham vs HU <50% (C), >0.9000 for each comparison (D).

**Fig 2, Supplemental 2:**
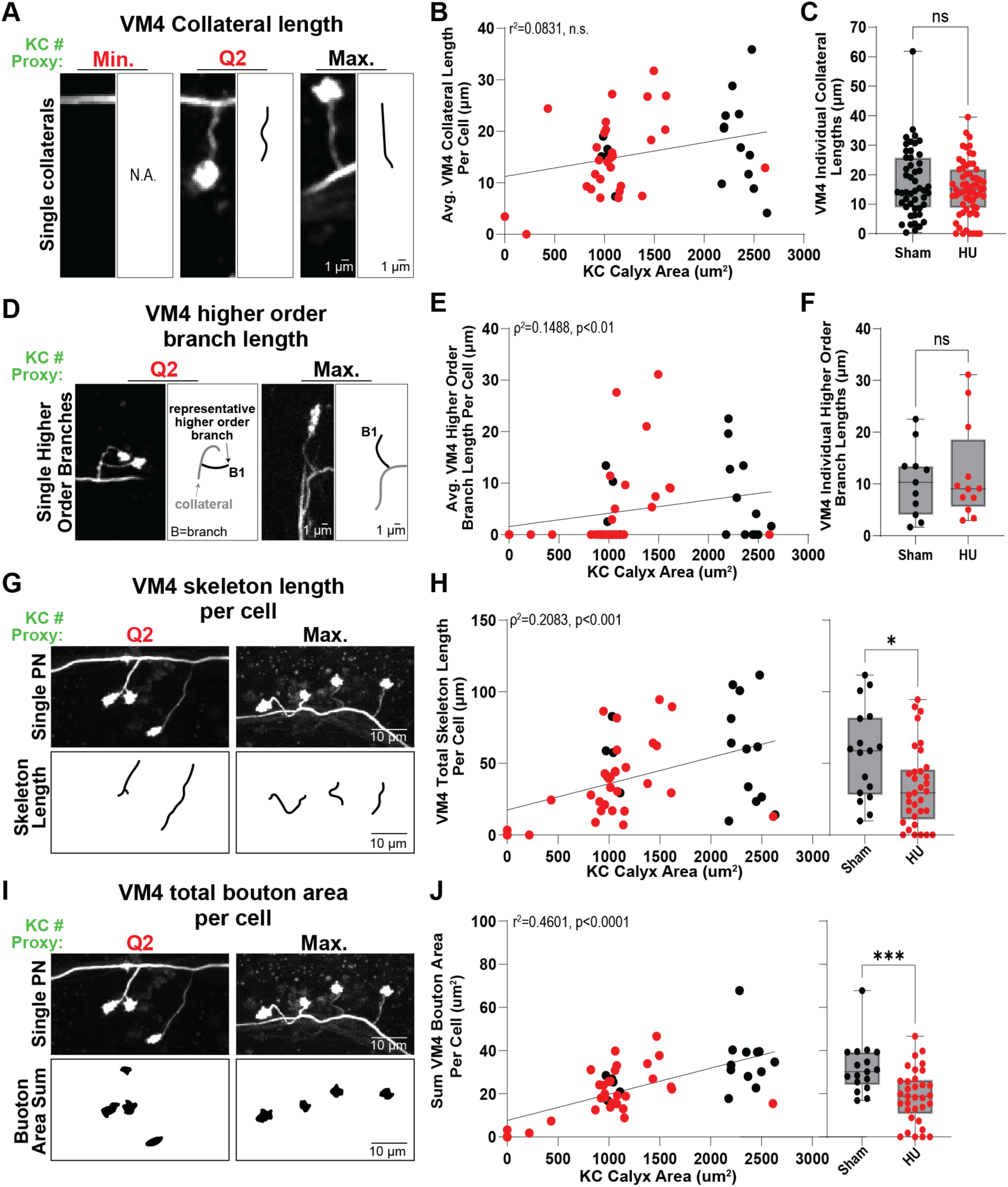
VM4 morphological features variably scale to KC population size. **A–C:** (A) Representative adult VM4 PNs showing differences in collateral growth with varying KC population sizes induced by HU ablation (red) or sham control (black). Cartoons depict traced collaterals. (B) Scatter plot showing the relationship between KC population size (proxy) and average collateral length per PN. Each point represents one cell. Statistics: simple linear regression; n.s. p-value: 0.052. (C) Box-and-whisker plot of collateral length across conditions. Each point represents a single collateral. Statistics: two-tailed Mann–Whitney test; n.s. p-value: 0.5269. **D–F:** (D) Representative higher-order branches from adult VM4 PNs. Cartoons depict traced branches. (E) Scatter plot of KC population size versus average higher-order branch length per PN. Each point represents a single cell. Statistics: two-tailed Spearman’s correlation; best-fit lines from simple linear regression. (F) Box-and-whisker plot of higher-order branch length across conditions. Each point represents a single higher order branch. Statistics: two-tailed Mann–Whitney test; n.s. p-value: 0.9150. **G-H:** (G) Representative PN skeleton lengths (i.e. the sum of all collateral and higher-order branch lengths per cell) for VM4 PNs. Cartoons depict traced skeletons. (H) (Left) Scatter plot of KC calyx area versus total VM4 skeleton length. Each point represents a single cell. Statistics: two-tailed Spearman’s correlation; best-fit lines from simple linear regression. (Right) Box-and-whisker plot of total skeleton length across conditions. Each point represents a single cell. Statistics: two-tailed Mann-Whitney test. \**p* < 0.05. **I-J:** (I) Representative VM4 PNs showing bouton area sums per cell across KC population sizes. Cartoons depict traced bouton areas. (J) (Left) Scatter plot showing clone calyx area versus bouton area sum per VM4 PN. Each point represents a single cell. Statistics: simple linear regression. (Right) Box-and-whisker plot of total bouton area per VM4 PN. Each point represents a single cell. Statistics: two-tailed Mann-Whitney test. \*\*\**p* < 0.001. Min.: Minimum. Q2: Quartile 2. Max.: Maximum.

**Fig 2, Supplemental 3:**
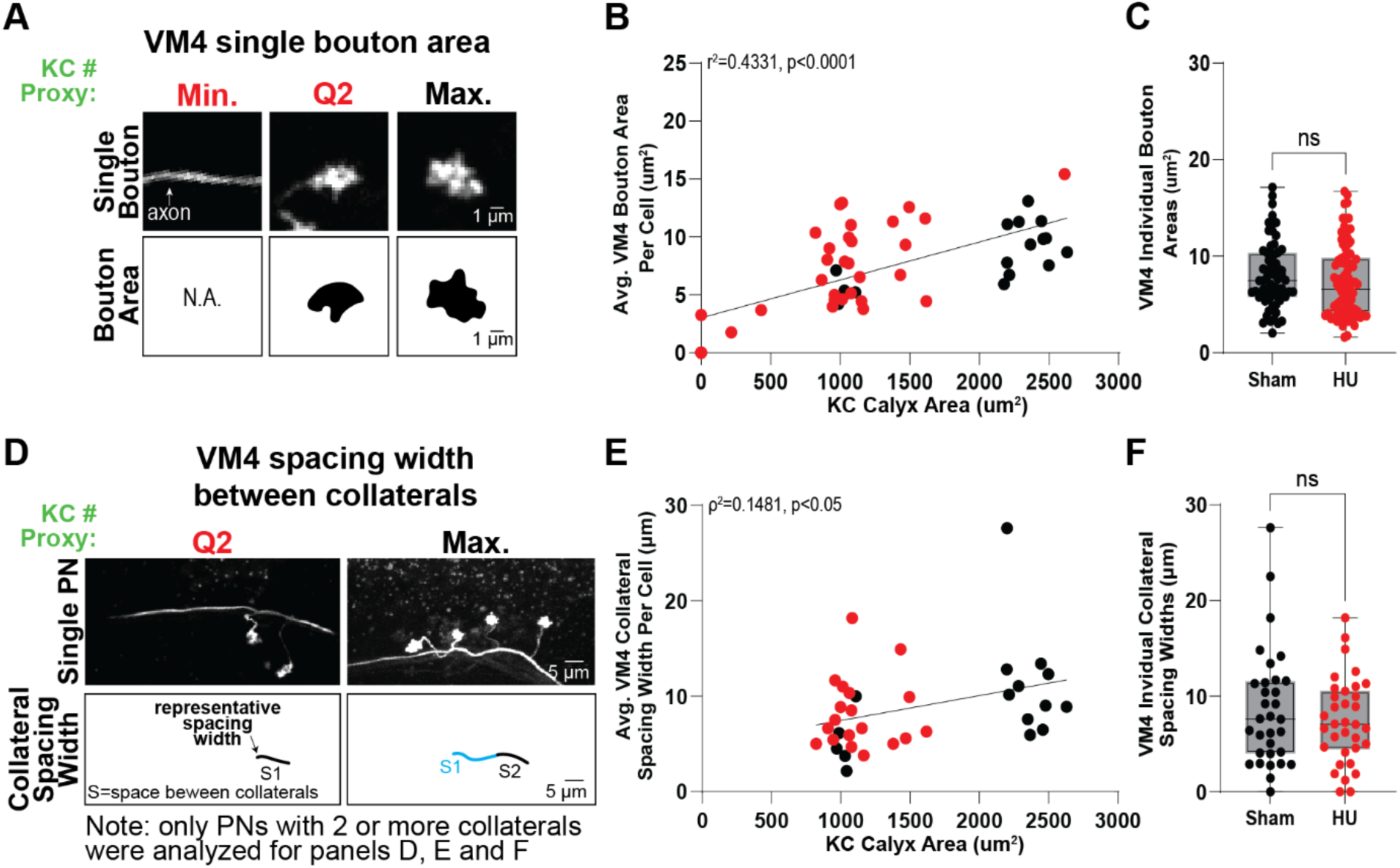
VM4 bouton size and collateral spacing show limited sensitivity to KC number. **A–C:** (A) Relationship between KC population size and individual bouton area for VM4 PNs across varying KC population sizes induced via HU ablation (red text), or sham control (black). Cartoons depict traced bouton areas. (B) Scatter plot showing the relationship between KC population size proxy and average bouton area for VM4 PNs. Each data point is a single cell. Statistics: simple linear regression. (C) Box-and-whisker plot describing measures of bouton area for VM4 PNs across control (black dots) and HU (red dots) conditions. Each data point is a single bouton. Statistics: two-tailed Mann-Whitney test. n.s. p-value: 0.2134. **D–F:** (D) Relationship between KC population size and spacing width between individual collaterals along VM4 PN axons across HU (red) and control (black) conditions. Cartoons depict traced collateral spacing. (E) Scatter plot showing the relationship between KC population size proxy and average collateral spacing width per PN. Each point represents one cell. Statistics: two-tailed Spearman’s correlation; best-fit lines from simple linear regression. (F) Box-and-whisker plot of individual collateral spacing width for VM4 PNs across control (black) and HU (red) conditions. Each point represents one spacing measurement. Statistics: two-tailed Mann–Whitney test; n.s. p-value: 0.6168. Min.: Minimum. Q2: Quartile 2. Max.: Maximum.

**Fig 3, Supplemental 1:**
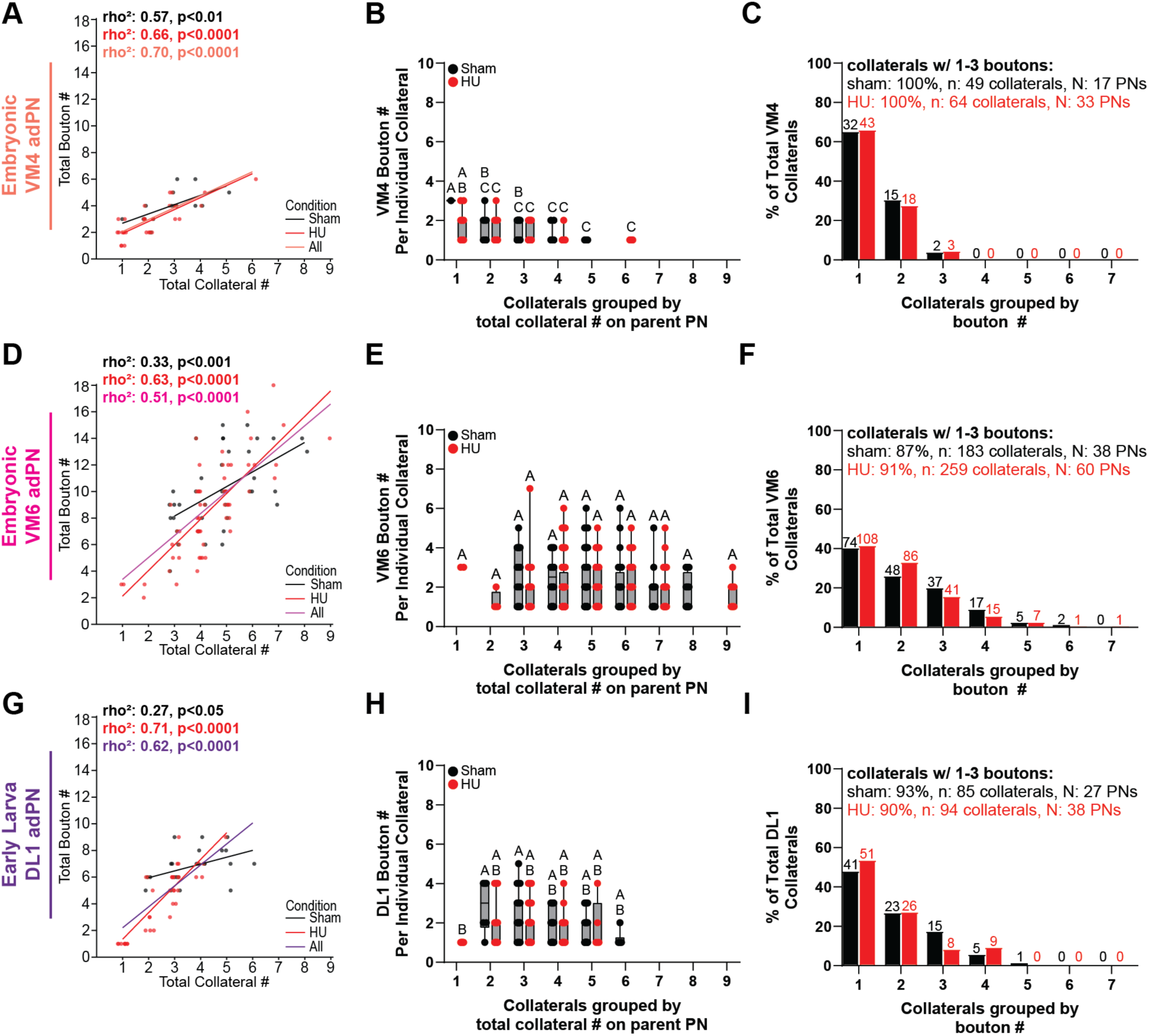
PN subtypes exhibit bouton–collateral modularity independent of KC population size. **A–C:** Single VM4 adPNs from HU experiments. (A) Scatter plot showing the relationship between total collateral number and total bouton number per PN; points colored by experimental condition as described in Figs. 1–2 (sham: black, HU: red). Scatter plots shown with jitter to reduce point overlap. (B) Box-and-whisker plots of bouton number per collateral, binned by the total collateral count of the parent PN. (C) Histogram showing the frequency distribution of bouton numbers per collateral. **D–F:** Single VM6 adPNs from HU experiments. Same plot types as A–C. **G–I:** Single DL1 adPNs from HU experiments. Same plot types as A–C. Statistics: (Scatter plots) Spearman’s correlation; best-fit lines generated by simple linear regression; each point represents one PN. (Boxplots) two-way ANOVA followed by Tukey’s post-hoc test; Compact Display System (CDS) letters indicate significant group differences (see Figure 3 supplemental table for details); each point represents one collateral.

**Fig 3, Supplemental 2:**
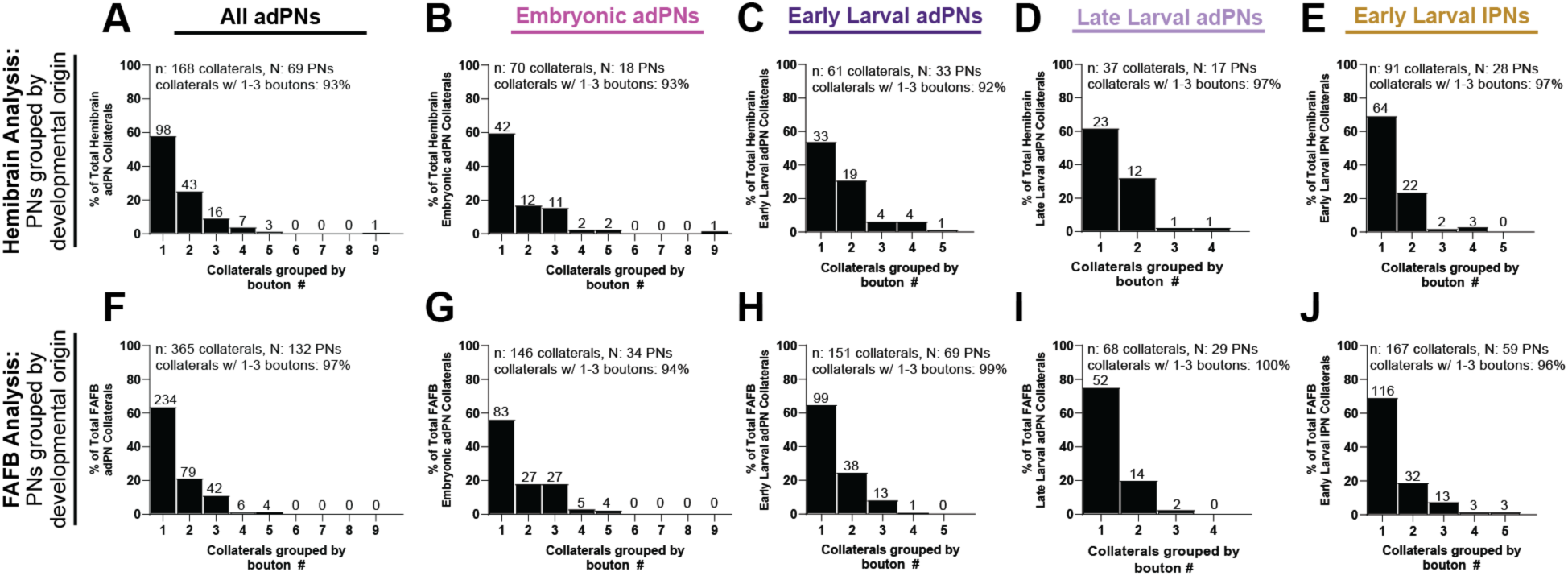
Discrete bouton number per collateral across PN developmental origins. **A–J:** Histograms showing the frequency distribution of bouton counts per collateral for PNs annotated in the hemibrain (A-E) or the FAFB/FlyWire (F-J) connectomes. (A,F) All PNs combined. (B,G) Embryonic adPNs. (C,H) Early larval adPNs. (D,I) Late larval adPNs. (E,J) Early larval lateral PNs (lPNs).

**Fig 5, Supplemental 1:**
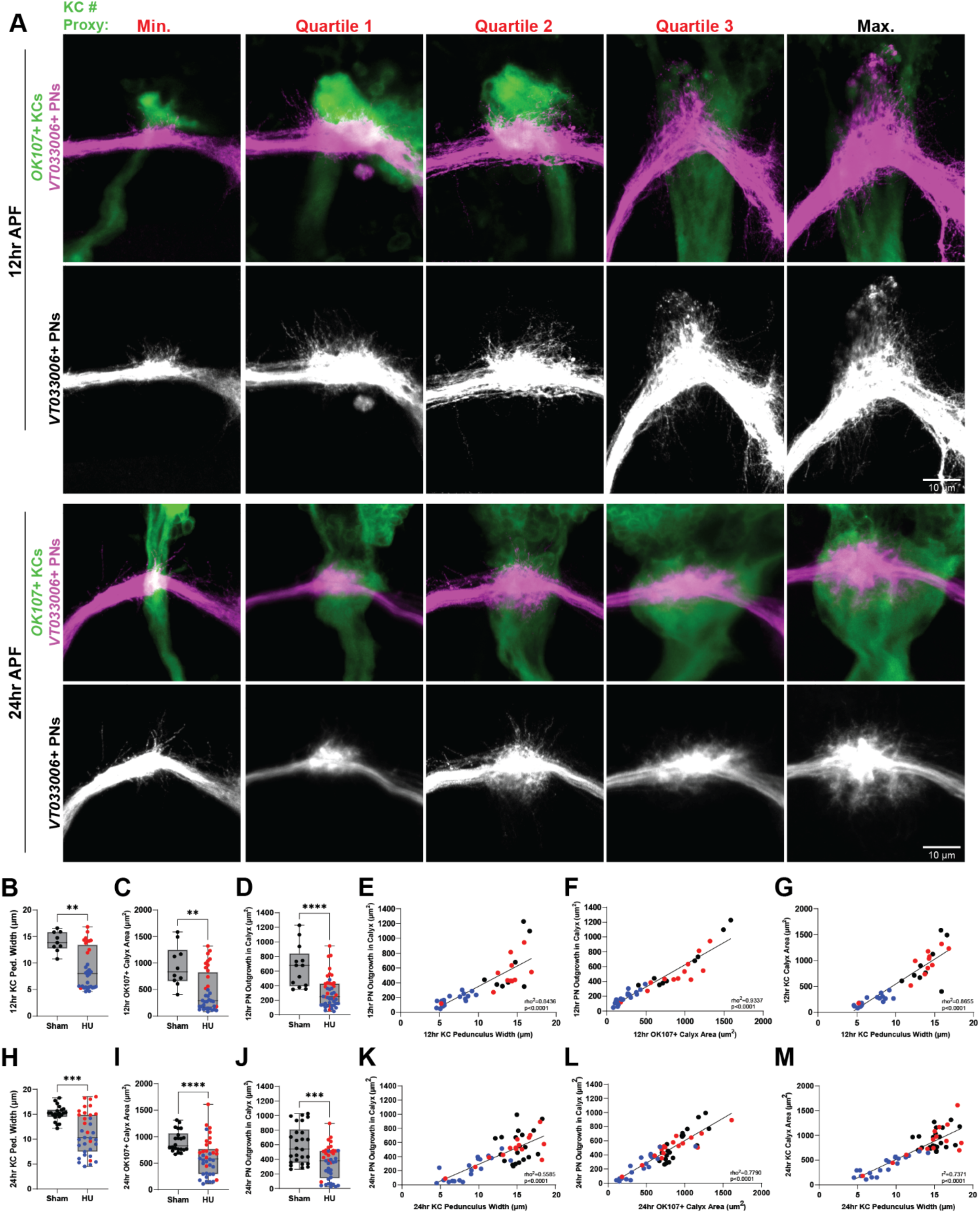
Early expansion of nascent collateral–bouton assemblages within the calyx is tuned by KC number. **A:** Representative images of *OK107*+ KCs (green) and *VT033006*+ PNs (magenta or gray) at 12hr APF (rows 1 and 2) and 24hr APF (rows 3 and 4) across varying KC population sizes induced via HU ablation (red titles), or sham treatment as control (black title). **B-D**: Box-and-whisker plots quantifying KC pedunculus width (B), OK107+ calyx area (C), or PN outgrowth in the calyx (D) at 12hr APF. Statistics: two-tailed Mann–Whitney test; ****p < 0.0001, **p<0.01. In panels B through M, Black points: Sham condition. Red and blue points, HU ablation; blue points are hemispheres in which ALl1 was lost, red points hemispheres in which it was retained. **E-F**: Scatter plots showing the relationship between KC pedunculus width and PN outgrowth (E), *OK107*+ calyx area and PN outgrowth (F), or KC pedunculus width and KC calyx area (G) at 12hr APF. Statistics: two-tailed Spearman’s correlation. Best-fit lines from simple linear regression. **H-J**: Box-and-whisker plots quantifying KC pedunculus width (B), OK107+ calyx area (C), or PN outgrowth in the calyx (D) at 24hr APF. Statistics: two-tailed Mann–Whitney test; ****p < 0.0001, ***p<0.001. **K-L**: Scatter plots showing the relationship between KC pedunculus width and PN outgrowth (E), *OK107*+ calyx area and PN outgrowth (F), or KC pedunculus width and KC calyx area (G) at 24hr APF. Statistics: two-tailed Spearman’s correlation (K, L) or simple linear regression (M). Best-fit lines from simple linear regression.

## Notes

### Competing Interest Statement

The authors have declared no competing interest.

